# Multiple LPA3 receptor agonist binding sites evidenced under docking and functional studies

**DOI:** 10.1101/2024.09.03.611107

**Authors:** K. Helivier Solís, M. Teresa Romero-Ávila, Ruth Rincón-Heredia, José Correa-Basurto, J. Adolfo García-Sáinz

**Author notes:** Correspondence: J. A. García-Sáinz. Inst. Fisiología Celular, UNAM Ciudad Universitaria, Ap. Postal 70-600 Ciudad de México 04510 México.

## Abstract

Comparative studies using (lysophosphatidic acid) LPA and the synthetic agonist, oleoyl-methoxy glycerophosphothionate, OMPT, in cells expressing the LPA3 receptor revealed differences in the action of these agents. The possibility that more than one recognition cavity might exist for these ligands in the LPA3 receptor was considered. We performed agonist docking studies exploring the whole protein to obtain tridimensional details of the ligand-receptor interaction. Functional in-cellulo experiments using mutants were also executed. Our work includes blind docking using the unrefined and refined proteins subjected to hot spot predictions. Distinct ligand protonation (charge-1 and-2) states were evaluated. One LPA recognition cavity is located near the lower surface of the receptor close to the cytoplasm (Lower Cavity). OMPT displayed an affinity for an additional identification cavity detected in the transmembrane and extracellular regions (Upper Cavity). Docking targeted to Trp102 favored binding of both ligands in the transmembrane domain near the extracellular areas (Upper Cavity), but the associating amino acids were not identical due to close sub-cavities. In the in-cellulo studies, LPA action was much less affected by the distinct mutations than that of OMPT (which was almost abolished). Docking and functional data suggest distinct agonist binding cavities in the LPA3 receptor

## 1. Introduction

Lysophosphatidic acid (LPA) is a bioactive lipid that exerts many of its actions through six G protein-coupled receptors (GPCRs) that constitute the LPA subfamily [1–3]. Among them is the LPA3 receptor, which mainly couples to Gαi/o and Gαq/11, participating in diverse signaling events that lead to various physiological and pathophysiological effects [4, 5]. Structurally, the LPA3 receptor is a membrane protein of 353 amino acids containing 7 transmembrane domains (TM), which are connected by three intracellular (IL) and three extracellular (EL) loops; it comprises an extracellular amino terminus and an intracellular carboxyl end [2, 4, 5]. Though it has not been crystallized yet, it has a similarity of approximately 53% with the LPA1 receptor, the best-studied member of this group [3, 6–8]. It is worth mentioning that the domains that regulate LPA1 receptor function are not conserved in the LPA3 receptor; these include a dileucine motif, a mono ubiquitination residue, and a PDZ pattern, among others [5, 9]. Structural analysis of the LPA3 receptor showed that it conserves motifs/domains typical of the A family of GPCRs, which comprise the CWxP, EHR, PIF, and NPxxY areas, each having a specific function during and after agonist attachment [2, 5, 10].

It is generally accepted that the receptor’s recognition cavities are flexible, and that ligand binding promotes conformational changes that impact their spatial configuration (https://pubmed.ncbi.nlm.nih.gov/33076289/). These dynamic events allow displacements of the intracellular loops and the TM regions, which promote G-protein recruitment and exposure of sites that can be phosphorylated by distinct protein kinases, including G protein-coupled receptor kinases (GRKs) and second messenger-activated protein kinases [10–13]. Receptor phosphorylation favors β-arrestin-receptor interaction, triggering its internalization and subsequent signaling events [14–16]. Determining how a ligand interrelates with a given receptor provides fundamental insights into the search for drugs that could effectively regulate its activity and the participating mechanisms. In this sense, molecular docking simulations have become a method that allows a better knowledge of the receptor regions in which the ligand could bind, suggesting the possible residues involved [17].

Pioneer structural work with the LPA3 receptor identified amino acids that interact with the polar head of LPA [7, 18–21]. When these residues were mutated, its function was altered, particularly those in the TM domains 3, 4, 5, and 7, mainly involving the following sites: Arg105 (3.28), Gln106 (3.29), Trp153 (4.64), Arg209 (5.38), and Lys296 (7.35) (in parentheses are the Ballesteros and Weinstein [22] numbering scheme; these numbers were obtained from the server: https://gpcrdb.org/protein/lpar3_human/ [23]. It is relevant that mutation of residues Trp153 (4.64) and Lys296 (7.35) directly impacts ligand binding and activation of the LPA3 receptor, i.e., LPA potency and efficacy were decreased in RH7777 cells transfected with mutant receptors when compared to those expressing the wild type [20].

No specific agonist for the LPA3 receptor subtype is available since most also activate the LPA1 and LPA2 receptors. A ligand that shows high potency for LPA3 is 1-oleoyl-2-methyl-sn-glycero-3-phosphothionate (OMPT) [24]. OMPT and related analogs exhibit affinity for different LPA receptors, but they have a weak selectivity for the LPA receptor subtypes [25–27]; despite these selectivity problems, OMPT remains a reference for studies on LPA3 receptors. To our knowledge, the binding site for OMPT in LPA receptors has not been explored.

It was recently reported that LPA and OMPT exhibit marked pharmacodynamic differences in LPA3-expressing cells [28]. In particular, two findings were astonishing: firstly, OMPT exerts its effects within two very distinct concentration ranges (1-10 nm and > 100 nM), and secondly, LPA induced a “receptor refractory state” that blocks calcium response to re-challenging with this natural agonist; however, under the same conditions, OMPT was able to produce a conspicuous response [28]. Both groups of data suggest the possibility that distinct binding cavities might exist for these agonists in the LPA3 receptor: one showing a very high affinity for OMPT and another one, likely shared by LPA and OMPT (which are structural analogs) with a lower affinity for both agonists in the range of 100-300 nM [28]. Intrigued by such findings and lacking structural information on the OMPT-LPA3 receptor interaction, we performed ligand-receptor docking studies to get insights into their protein binding pocket(s). The docking examination and the hot spot predictions indicate the presence of at least two distinct ligand recognition cavities, which might explain the observed functional responses to OMPT. In addition, receptor mutants were constructed and expressed, and their response to these agonists was studied. One of the mutants showed activity in reaction to LPA and essentially no effect on OMPT, which is consistent with the proposal of multiple binding pockets for these ligands [28]. This report documents distinct agonist identification cavities via docking simulations and in-cellulo experiments with receptor mutants. The findings open new experimental possibilities and might help better understand LPA3 receptor-ligand interaction and aid in developing new agonists and antagonists of potential therapeutic use.

## 2. Results

### 2.1 LPA3 receptor model without refinement

The LPA3 receptor has not been crystallized yet; therefore, its most likely 3D structure was obtained using different servers for configuration prediction (additional information is provided in section 4.1). Supplementary Table S1 enlists the values obtained using these servers and programs. Protein arrays from the distinct patterns were subjected to sequence alignments (Supplementary Fig. S1), and the Swiss Model was preferred because the best structural score values were obtained, and the resulting sequence was identical to the UniProt Q9UBY5 entry.

The unrefined 3D pattern from the Swiss Model program was employed to determine the binding parameters (free energy values (ΔG), Kd, and residue interaction) for the LPA3 receptor agonists, LPA and OMPT (a 126 Å3 grid box centered on the protein with (x, y, and z) coordinates in grid center for both agonists: 6.014,-0.124, and-2.385). Molecular docking simulation showed that the LPA3 receptor recognition cavities for LPA and OMPT were different (Supplementary Table S2), and the binding energy value was lower for OMPT than for LPA. The residues participating in ligand-receptor interaction are also indicated in Supplementary Table S2; please notice that the amino acids involved are listed in the order indicated by the program, considering their importance in the ligand-receptor interaction. Supplementary Fig. S2 depicts such interrelations using the unrefined model. OMPT (red) associates with amino acids located in TM regions 2 and 7 and the EL 1, 2 (we name this broad region Upper Cavity; see Supplementary Fig S2), whereas LPA (blue) reached a distinct site (broad region named Lower Cavity) of the LPA3 receptor structure with molecules preferentially located in IL 2 and IL 3, very close to the cytoplasm (Supplementary Fig. S2). The cartoon shown in Supplementary Fig. S3 depicts the LPA-interacting residues (blue) and OMPT-associating amino acids (red) (notice the remarkably different distribution of such binding cavities).

The 2D maps of the ligand-receptor interrelations obtained from the Discovery Studio program are also shown (Supplementary Figs. S4 (LPA) and S5 (OMPT)). This methodology allows adaptations of the ligand to obtain the best possible interaction with the receptor and performs the necessary adjustments to achieve the most favorable association; it also permits determining the agonist orientation, identifying the receptor’s amino acids involved, and defining the type of interaction (Supplementary Table S2). Notice that the phosphate group of LPA is fully protonated (i.e., net charge 0; Supplementary Figure S4), but in contrast, the phosphorothioate group of OMPT is partially deprotonated (net charge-1; Supplementary Figure S5) when interacting with the distinct LPA3 receptor-ligand recognition cavities.

The polar head groups of agonists such as LPA and OMPT seem important when these ligands bind to LPA3 receptors and for their activation [7, 18–20, 25]. LPA’s phosphate and OMPT’s phosphorothionate can be protonated (net charge 0), partially deprotonated (net charge-1), or fully deprotonated (net charge-2) at distinct pH values. Experimentally defined pKa values for these compounds have not been reported, but those of the analog, phosphatidic acid, are available (pKa1=3.0 and pKa2=8.0; https://avantiresearch.com/tech-support/physical-properties/ionization-constants). Assuming that the pKa values of LPA and OMPT could be similar to those of phosphatidic acid, at a pH of 7.4, the protonated species (charge 0) would represent < 0.1% of the total amount of the compound, whereas the partially deprotonated (charge-1) and fully deprotonated (charge-2) species would represent ≈ 79% and ≈ 20% of the total, respectively. The docking simulations using the unrefined receptor structure employed the protonated agonist species (charge 0). For all the following simulations, the three molecular species were employed (i.e., species with net charges 0,-1, and-2).

### 2.2 LPA3 refined Model

The molecular docking studies of protonated (charge 0) LPA (Fig. 1; blue) show that it occupied the Lower Site of the receptor, interacting with the IL 2 and 3, which, like in the LPA3 unrefined model, was very close to the cytoplasm. On the other hand, OMPT (charge 0) (Fig. 1; red) associates with the amino acids located in the Upper cavity, including TM 5, 6, 7, and 8, as well as the extracellular loop 3 (Fig. 1). Interestingly, simulation with both charge-1 ligands resulted in inversion of the agonist position in these cavities (Fig. 1). Simulation with charge-2 derived in interaction patterns similar to those observed with the uncharged ligands (Fig. 1). Tables 1 (for LPA) and 2 (for OMPT) provide the ΔG and Kd values and the list of interacting amino acids defined for the distinct conditions. Simulation with LPA (charge-2) resulted in a more negative ΔG and lower Kd values compared to the other conditions (Table 1); in contrast, modification of OMPT charge had only a minor impact on these parameters (Table 2). Cartoons showing the interrelating amino acids under these conditions are presented in Supplementary Fig. S6, and the 2D schemes showing the interacting residues and the type of association are presented in Supplementary Figs. S7.

**Figure 1.**
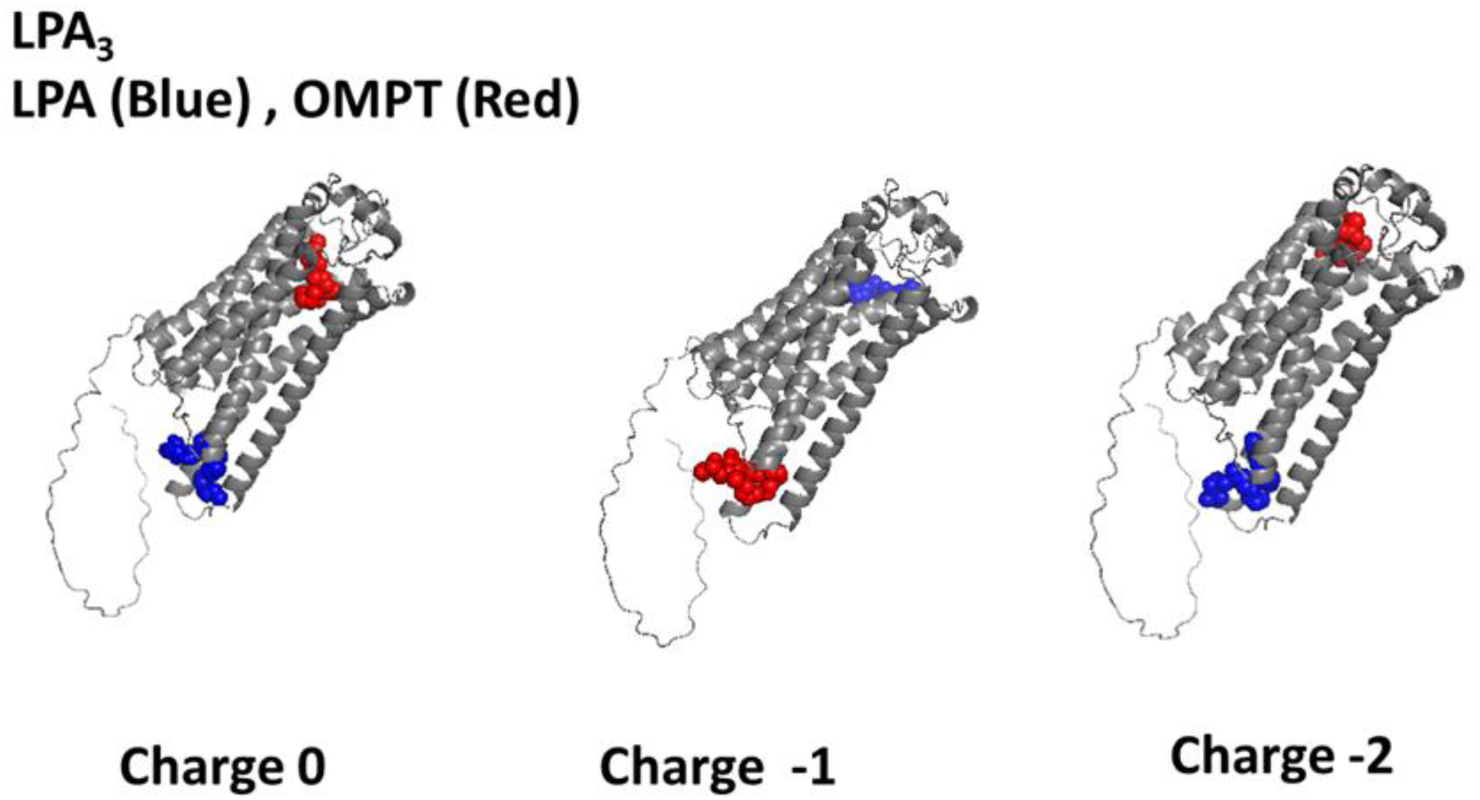
3D images resulting from the LPA3-LPA (blue) and LPA3-OMPT (red) docking analysis using PyMol. The refined structure was employed. Ligand net charges are indicated.

**Table 1.** Thermodynamic parameters and interacting amino acids were determined in the distinct docking simulations using LPA as the ligand.

| **Focus**<br>**(Charge)** | **$\Delta G$**<br>**(Kcal/mol)** | **Kd**<br>**($\mu M$)** | **Interacting**<br>**Amino acids** |
| --- | --- | --- | --- |
| **NONE**<br>**(0)** | -5.01 | $\approx 210$ | Val219, Leu220, Ser221,<br>Lys216, Arg230, Pro234,<br>Val136 |
| **NONE**<br>**(-1)** | -4.10 | $\approx 995$ | Arg276, Trp277, Val22,<br>His273, Val274, Lys 29,<br>Trp25, Ile 32, Leu 280,<br>Val33, Val 36, Tyr83, Gly<br>37, Leu283, Met 87, Leu86,<br>Cys41 |
| **NONE**<br>**(-2)** | -7.18 | $\approx 5$ | Lys216, Thr217, Ile131,<br>Met134, Arg230, Pro234,<br>Leu237, Arg127, Val136 |
| **Trp102**<br>**(0)** | -5.99 | $\approx 40$ | Leu283, Leu280, Leu279,<br>Met87, Val33, Thr90, Trp25,<br>Arg276, Lys29 |
| **Trp102**<br>**(-1)** | -4.10 | $\approx 990$ | Gln106, Arg105, Lys275,<br>Leu86, Tyr14, Leu109,<br>Trp102, Ala180, Leu279,<br>Thr90, Asp23, Gly91, |
|  |  |  | Arg276, Val36, Val33,<br>Leu280, Trp277, Trp225,<br>Lys29, Ile32 |
| **Trp102**<br>**(-2)** | -8.93 | $\approx 0.30$ | Trp277, Leu279, Arg276,<br>Gly37, Val33, Trp25, Lys29,<br>Thr90, Leu86, Met87 |
| **Tyr293**<br>**(0)** | -5.79 | $\approx 60$ | Leu220, Lys216, Thr217,<br>Ile131, Val213, Pro234,<br>Arg135, Met134, Val136,<br>Asn139, His137, Phe63 |
| **Tyr293**<br>**(-1)** | -2.88 | $\approx 8000$ | Thr146, Lys142, Val145,<br>Trp153, Ile149, Leu150,<br>Tyr67, Val55, Ile56, His62,<br>Ala70, Asn71 |
| **Tyr293**<br>**(-2)** | -7.62 | $\approx 3$ | Trp277, Leu280, Leu279,<br>Leu283, Tyr83, Arg276,<br>Gly37, Val33, Trp25, Leu86,<br>Met87 |

**Table 2.** Thermodynamic parameters and interacting amino acids were determined in the distinct docking simulations using OMPT as the ligand.

| **Focus**<br>**(Charge)** | **$\Delta G$**<br>**(Kcal/mol)** | **Kd**<br>**($\mu M$)** | **Interacting**<br>**Amino acids** |
| --- | --- | --- | --- |
| **NONE**<br>**(0)** | -5.67 | $\approx 70$ | Trp277, Leu279,<br>Leu280, Leu283,<br>Arg276, Tyr14, Asp23,<br>Trp25, Ser94, Thr90,<br>Arg105, Leu86, Val33 |
| **NONE**<br>**(-1)** | -5.36 | $\approx 120$ | Lys216, Val213,<br>Ile131, Met134,<br>Arg230, Val136,<br>Arg135 |
| **NONE**<br>**(-2)** | -5.85 | $\approx 50$ | Ser94, Arg105, Leu86,<br>Asp23, Lys275,<br>Arg276, Leu279,<br>Val36, Trp277, Val33,<br>Ile32 |
| **Trp102**<br>**(0)** | -5.15 | $\approx 170$ | Leu283, Leu280,<br>Leu279, Tyr83, Val36,<br>Met87, Val33, Thr90,<br>Trp25, Arg276,<br>Trp277, Leu86 |
| **Trp102**<br>**(-1)** | -7.75 | $\approx 6$ | Leu256, Leu159,<br>Leu188, Phe278,<br>Lys275, Trp191,<br>Leu179, Ala180,<br>Tyr183, Tyr187,<br>Gln106, Arg105,<br>asp110, Ala282,<br>Leu279, Trp252 |
| **Trp 102**<br>**(-2)** | -7.11 | $\approx 6$ | Leu280, Leu279,<br>Leu283, Ile32, Gly37,<br>Val36, Val33, Arg276,<br>Trp25, Met87 |
| **Tyr293**<br>**(0)** | -4.64 | $\approx 395$ | Arg230, Leu220,<br>Lys216, Thr217,<br>Arg231, Ile131,<br>His137, Val136,<br>Thr233, Pro234 |
| **Tyr293**<br>**(-1)** | -5.76 | $\approx 60$ | Arg232, Met235,<br>Lys239, Lys236,<br>Thr240, Tyr295,<br>Glu298, Ser294,<br>Tyr301, Ile291 |
| **Tyr293**<br>**(-2)** | -5.64 | $\approx 75$ | Lys296, Lys236,<br>Thr240, Lys305,<br>Ile308, Tyr295,<br>Glu298, Tyr301,<br>Ser294, Met304, Ile291 |

### 2.3 Trp102-focused model

Trp102 is located near the Upper Cavity or the LPA3 receptor configuration (Supplementary Fig. 8). As previously mentioned, according to 3D structural analysis with PyMol., this residue was consistently found in the hot spot predictions and visualized as exposed and centered in a cavity. Trp102-focused docking was performed for LPA and OMPT with distinct ligand protonations as indicated. Independently of the molecule charge, both agonists interacted with the receptor’s Upper cavity (Fig. 2). The uncharged ligands interrelated with the TM regions 1, 2, and 7. LPA (charge-1) interconnects with TM domains 1, 2, 3, and 7 and EL2, whereas OMPT (charge-1) binds to amino acids in TM domains 3, 5, 6, and 7 and EL 3. Simulations with both charge-2 ligands lead to their interaction with residues in TM 1 and 7 and EL 2. Cartoons showing the participating amino acids under these conditions are presented in Supplementary Fig. S9, and the 2D schemes showing the interacting residues and their type of association are illustrated in Supplementary Fig. S10. For LPA (Charge-2), Trp102-focused docking resulted in a more negative ΔG and lower Kd values (Table 1); similarly, OMPT (charges-1 and-2) also derived more favorable thermodynamic values (Table 2). The cartoons and the 2D images (Figs. S9 and S10) evidenced that both ligands shared some amino acids under these conditions.

**Figure 2.**
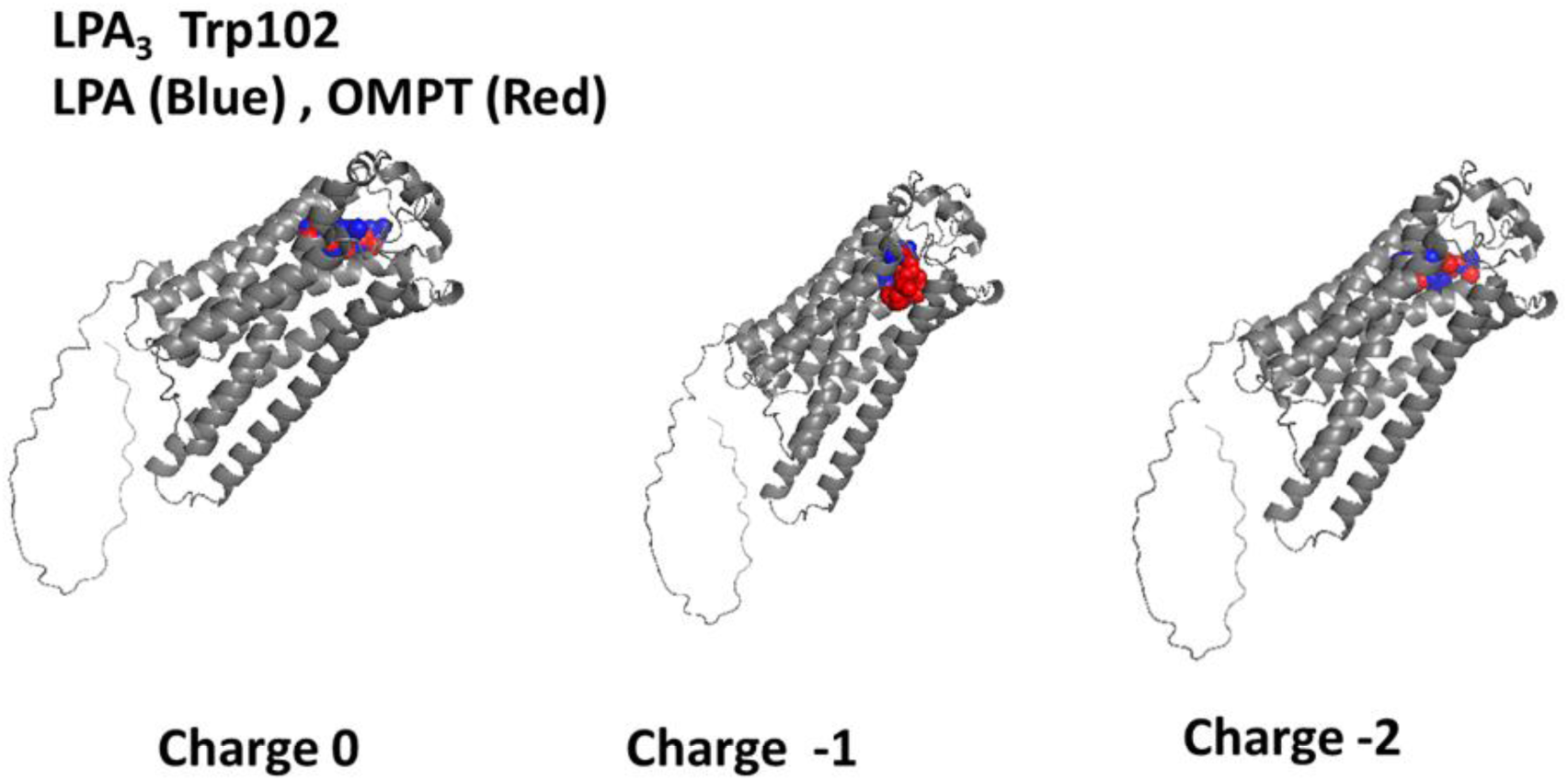
Images resulting from the docking analysis of the refined 3D structure of LPA3 receptor (Trp102 focused) interacting with LPA (blue) and OMPT (red). Ligand net charges are indicated.

### 2.4 Tyr293-focused model

The localization of Tyr293 in the LPA3 structure is shown in Supplementary Fig. S8. This residue participates in the attachment of the LPA antagonist, Kil6425, and an extensive series of pharmacological agents used in therapeutics [29]. Figure 3 presents the non-bond interaction of LPA (blue) and OMPT (red) with the LPA3 receptor when the docking analysis is focused on Tyr293. Using ligands with charge 0 for the docking simulation showed that the LPA binding cavities were located near the cytoplasm in the intracellular loops 1, 2, and 3. OMPT-LPA3 receptor interaction occurred in the Upper Cavity involving TM domains 1, 7, and EL 2 and 3. When simulations were performed using both ligands with charge-1, the agonists interrelated with the lower part of the receptor (LPA) IL1, IL2, and TM 4; for OMPT binding residues at IL3, Helix 8 and TM 7 played a role. With the receptor focused on Tyr 293, both molecules with charge-2 interacted with a pattern opposite to that observed at charge 0. Cartoons illustrating the partaking amino acids under these conditions are presented in Supplementary Fig S11, and the 2D schemes showing the participating residues and their type of association are depicted in Supplementary Fig12. Table 1 shows that LPA (charge-2) exhibited a relevant negative ΔG and a small Kd value. OMPT charge causes minor modifications of these thermodynamic parameters.

**Figure 3.**
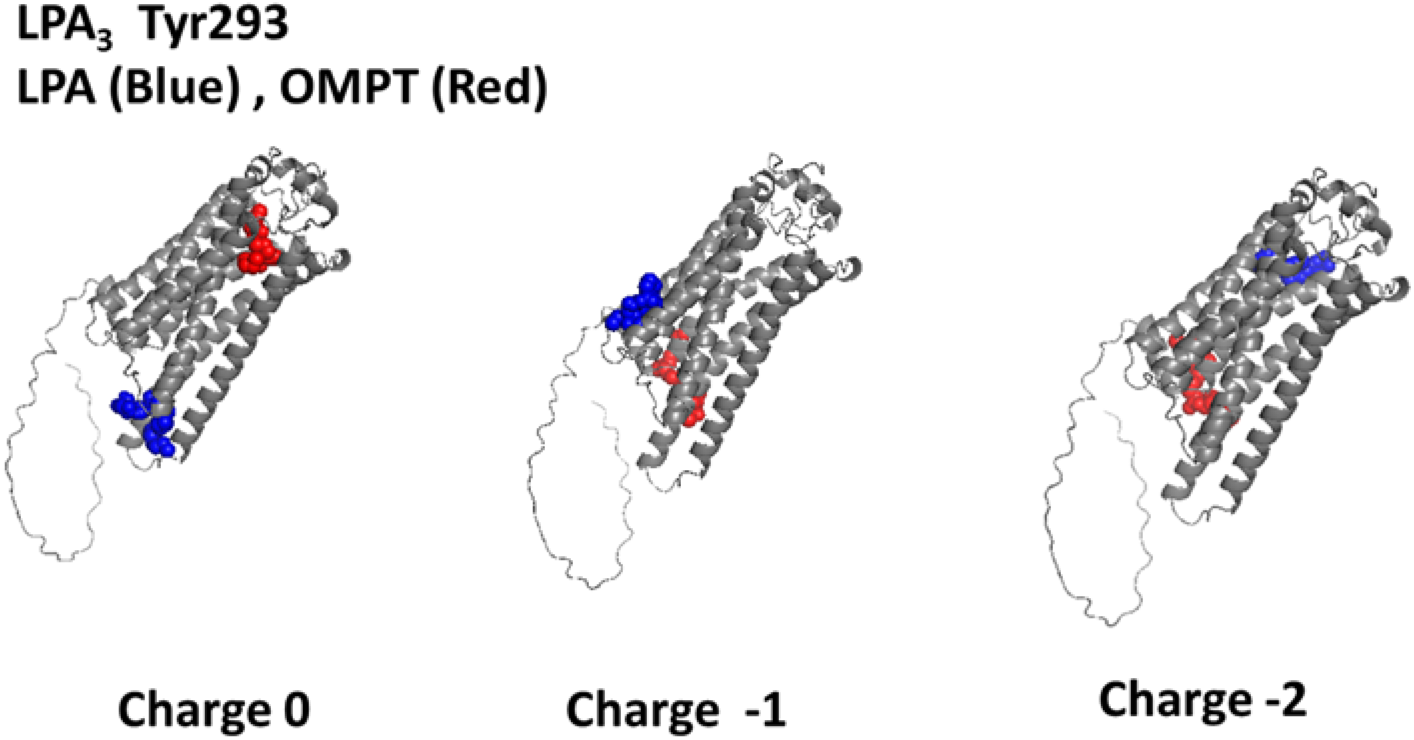
Images resulting from the docking analysis of the refined 3D structure of the LPA_3_ receptor (Tyr293 focused) interacting with LPA (blue) and OMPT (red). Ligand net charges are indicated.

### 2.5 In cellulo signaling experiments

Mutant receptors in which the ligand-interacting residues (unrefined model) in the cavities were substituted for alanine were expressed, and their calcium response to 1 µM LPA and 1µM OMPT was determined. The reaction of cells manifesting the wild-type receptor was considered the reference and was compared to the mutants in the upper cavity, lower cavity, and both.

Representative calcium tracings were presented in Fig. 4, and the data analysis is shown in Fig. 5. In the control cells, as anticipated, 1µM LPA or 1µM OMPT triggered an almost immediate increase in intracellular calcium. In contrast, a very weak response to OMPT was observed with the distinct mutants. The reaction to LPA was diminished but not blocked, and the Upper Cavity mutant showed a robust effect. These data indicate that some residues are much more relevant for the action of OMPT than for that of LPA. Similar findings were obtained with these mutants when ERK 1/2 phosphorylation was studied (i.e., the response to LPA was diminished in cells expressing the mutant receptors, while that to OMPT was almost abolished) (unpublished data).

**Figure 4.**
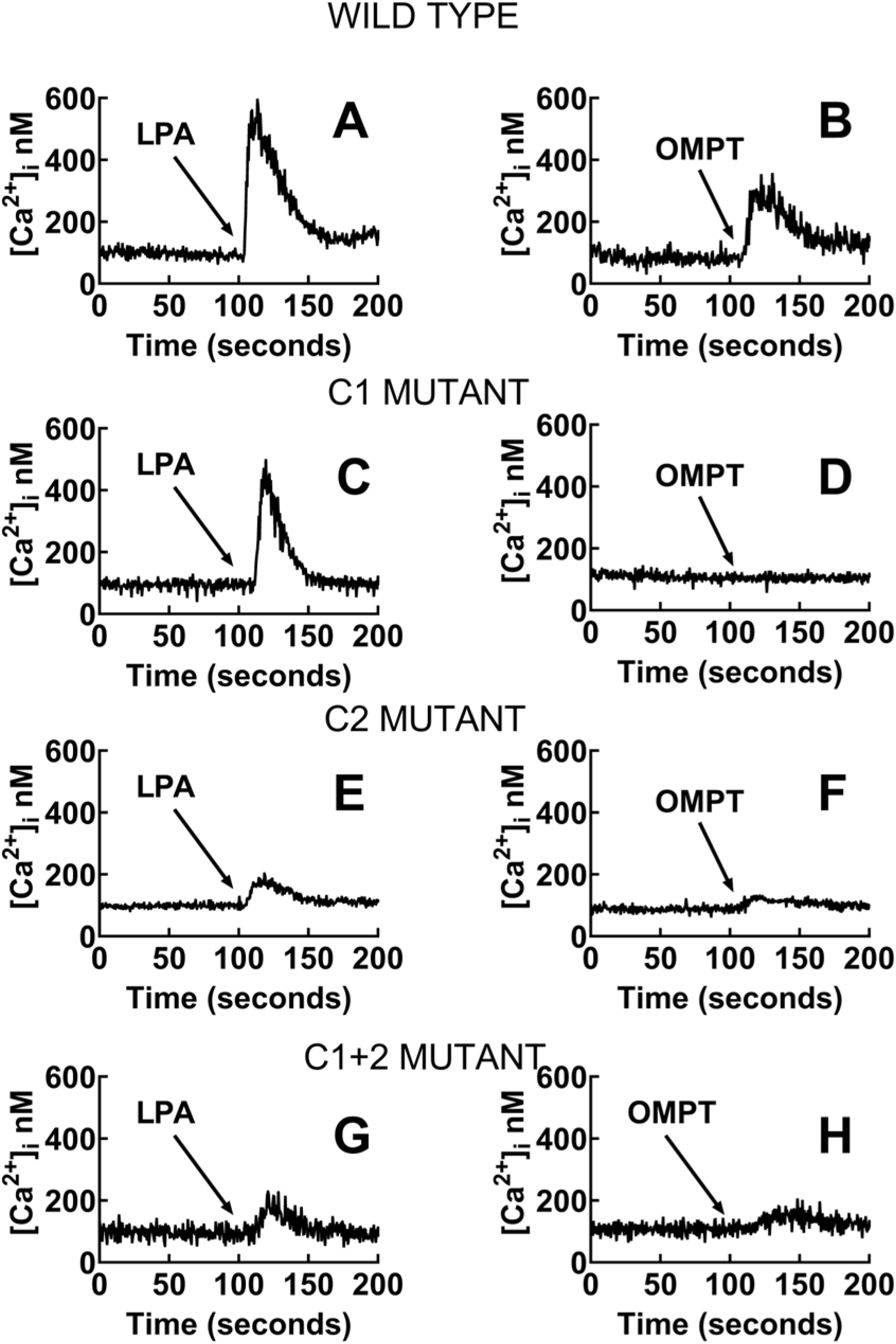
Representative calcium tracings were obtained in cells expressing the wild-type LPA_3_ receptors or mutants with substitutions in Cavity 1 (C1 mutant), Cavity 2 (C2 mutant), or both Cavities (C1+2 mutant). Cells were challenged with 1 µM LPA (right) or 1 µM OMPT (left).

**Figure 5.**
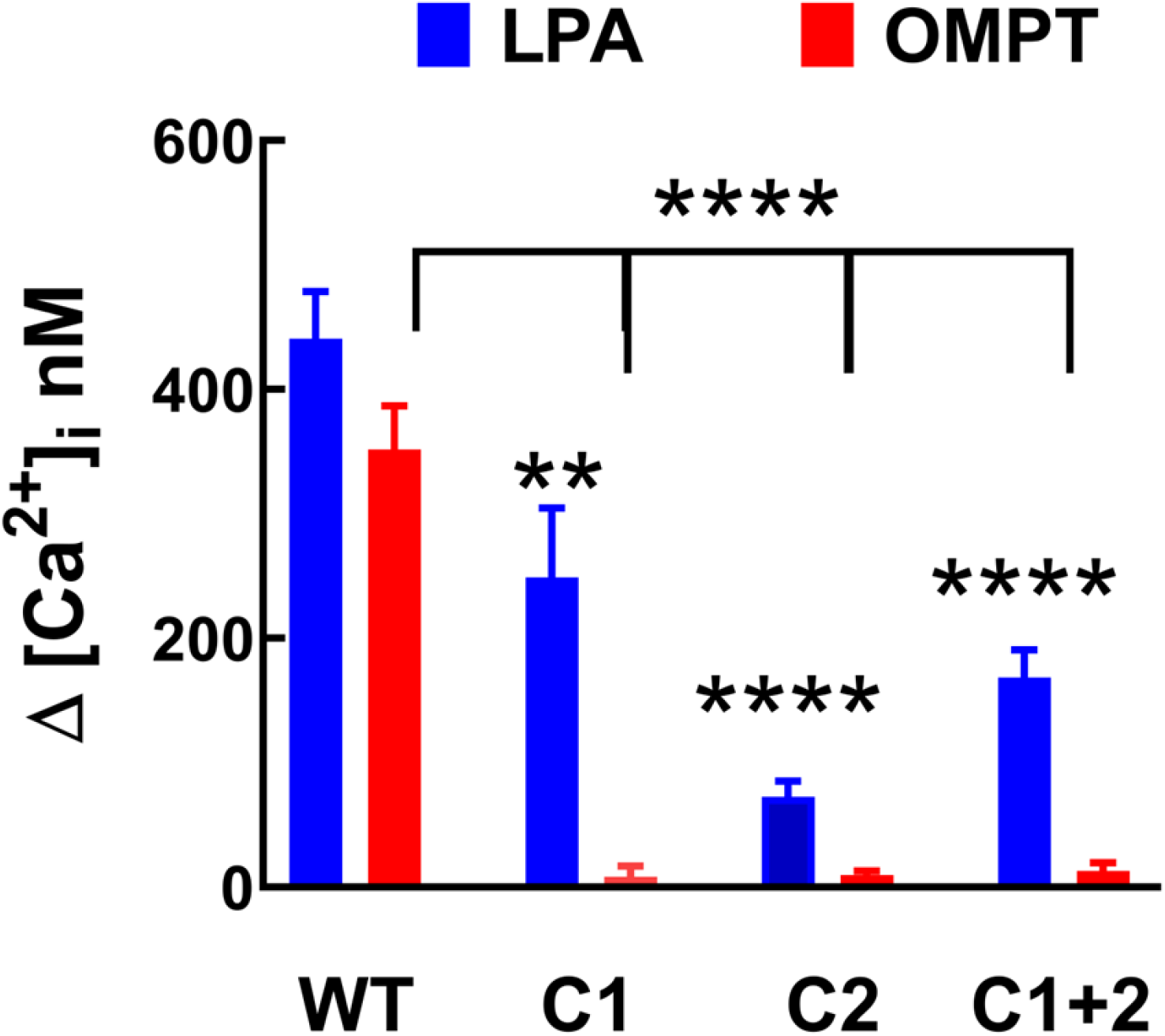
Calcium tracings were obtained in cells expressing the wild-type LPA_3_ receptors or mutants with substitutions in Cavity 1 (C1 mutant), Cavity 2 (C2 mutant), or both Cavities (C1+2 mutant). Cells were challenged with 1 µM LPA or 1 µM OMPT. The means are plotted, and vertical lines indicate the SEM of 4-5 experiments performed on different days using distinct cell cultures. **p < 0.01, and ****p < 0.0001 vs. cell expressing the wild-type receptor

## 3. Discussion

The present work was designed to define if LPA and OMPT bind to a single cavity or different agonist recognition pockets in the LPA3 receptor. The possible existence of more than one ligand-identification region was suggested in experiments using cultured cells [28]. Blind docking studies were performed using the predicted configuration of LPA3 (UniProt, Q9UBY5 entry). 3D structures of refined and focused (to Tyr293 and Trp102) docking analyses were carried out employing the distinct protonation states of the ligands. As indicated, Tyr293 was identified as a critical amino acid for several agents interacting with the LPA3 receptor [29], and Trp102 was consistently detected in the hot spot prediction.

In our work, we observed the docking of LPA and OMPT to two principal LPA3 receptor regions named, for this work, Upper and Lower Cavities. This pattern was noticed in both the unrefined and refined docking analysis and with changes in the net charge of the ligands. In fact, under most of these conditions LPA was localized in the Lower Cavity. This location was reversed when a charge of-1 was assigned to both ligands. Previous work [7, 18–21] has detected a central binding cavity with emplacement similar to the Upper Cavity reported here. In these studies, a box of 60 Å3 was employed, whereas in our work, we used a 126 Å3 box and a grid space of 0.375 Å3. Therefore, it seems likely that space limitations (i.e., using a box of 60 Å3) might have impeded the Lower Cavity detection. Avoiding expanse restrictions has positive and negative aspects. When the ligand recognition cavity(s) is not known, it seems convenient to explore the entire protein surface by docking, a procedure named “blind docking”, which implies using a large box [30]; however, focusing on predicted binding regions is known to improve docking accuracy and efficiency [30].

It should be mentioned that the Upper Cavity is in the TM core of the receptor, where most agonists associate with class A GPCRs. The Lower Cavity observed in this work included ligand interaction with ICLs and hardly any contact with the TM domains, which is an uncommon feature. This Lower Cavity might represent a gating point for the molecules rather than an agonist-activating region. However, with the current information, determining the functional relevance of these Cavities is impossible. The operational data obtained with the mutants of these pockets indicate that amino acid substitution in any of them markedly altered the action of OMPT but, to a much lesser extent, that of LPA. These findings support the idea that the effect of these two agonists does not involve the same receptor associations but rather that distinct residues participate in receptor activation.

Previous work, also using docking/ computational modeling and mutants, identified that the LPA3 receptor interacted mainly with the polar head of LPA at Arg105 (3.28), Gln106 (3.29), Trp153 (4.50), Arg209 (5.60), Arg276 (7.36) and Lys275 (7.35) [7, 18–21]. These amino acids were considered critical for LPA binding to the receptor and their mutation decreased agonist efficacy and potency and as expected, such modifications eliminated ionic interactions in the modeled LPA3-ligand prototype [7, 18–21]. It is important to mention that the same amino acids were identified in our docking studies. The residues detected were Arg276 (9 times), Arg105 (4 times), Lys275 (3 times), Gln106 (2 times), and Trp153 (1 time); this information includes data for LPA and OMPT. Other interacting amino acids frequently found in the docking simulations for both ligands include: Val33 (9 times), Trp25 (7 times), Trp277 (7 times), Leu280 (7 times), Leu86 (7 times), and Leu283 (6 times) among others.

The similarities of our present findings with those published earlier [7, 18–21] suggest the possibility that the LPA binding site observed in the Upper Cavity might correspond to that previously determined [7, 18–21]. No prior docking data for LPA3-OMPT interaction has been published. Our findings indicate that OMPT occupies the same Upper Cavity as LPA. Interestingly, the functional information formerly published [28] and that using mutants presented here strongly suggest that differences among these agonist-recognizing regions exist; therefore, they should be considered distinct, although some interacting residues could be shared. Exploration of the LPA3 receptor cavities, using PyMol, showed many capable of ligand binding (Supplementary Fig. S13). The localization of the agonists was included in the analysis, employing the Trp102 docking simulations (charges-1 and-2). It can be observed that LPA and OMPT occupy close but not identical positions; in addition, it seems likely that molecules with distinct charges (i.e., mixtures of agonists with charges-1 and-2) might interact with LP3. Very recent data using single-particle cryo-EM structures of human S1P1 receptor bound to Siponimod and heterotrimeric Gi complexes, as well as human LPA1 receptor and Gi complexes in the presence of LPA indicated that the ligands adopt different conformations to interact with their cognate GPCRs [31]. Such distinct molecule configurations might participate in the agonist-induced LPA3 activation. Similar structural approaches likely would be required to define the precise residues that interact with the different ligands and their diverse conformations. The present findings open new research possibilities to explore this crucial step in LPA3 receptor signaling, considering the physiological and physiopathological events in which it is involved [4, 5].

## 4. Materials and Methods

### 4.1 Modeling of LPA3 receptor structure, Gene ID:23566

The LPA3 receptor has not yet been crystallized. Therefore, we were forced to predict its most likely 3D structure using the amino acid sequence obtained from the Uniprot database (Q9UBY5 entry) (https://www.uniprot.org/uniprotkb/Q9UBY5/entry). Three of the most frequently utilized servers for structure prediction were employed: SWISS-MODEL (https://swissmodel.expasy.org/), I-TASSER (https://zhanggroup.org/I-TASSER/), and AlphaFold database (https://alphafold.ebi.ac.uk/). Supplementary Table S1 enlists the values obtained using these servers and programs. Subsequently, protein arrays from the three models were subjected to sequence alignments (Supplementary Fig. S1) employing the PROMALS3D server (http://prodata.swmed.edu/promals3d/promals3d.php) to determine the gaps or residue insertions that improve the arrangement identity with the 3D protein templates during the folding of the molecule. Although the data obtained showed that the 3D construction built with AlphaFold was the best evaluated structurally, the alignment revealed that several mutations or deletions improved the 3D configuration quality, possibly eliminating residues that can interact with the ligands. Therefore, the Swiss Model was preferred because the best values were estimated by ERRAT and a second evaluation of Ramachandran maps; additionally, the sequence obtained was identical to the UniProt Q9UBY5 entry.

### 4.2 Refined and validation of the LPA3 receptor 3D structure

Protein 3D structure refinement was carried out to perform accurate docking simulations. The arrangement obtained from the Swiss Model server was subjected to molecular refinement using ReFOLD (https://www.reading.ac.uk/bioinf/ReFOLD/ReFOLD3_form.html). The refined protein with the best score was used for the docking study to explore receptor cavities capable of ligand recognition.

### 4.3 Search for ligand-binding cavities on the LPA3 receptor

The hot spot prediction was performed to identify possible ligand-recognition cavities using the BetaCavityWeb (http://voronoi.hanyang.ac.kr/betacavityweb/) and the Fpocketweb (http://durrantlab.com/fpocketweb) servers. The protein from the Swiss Model server, as well as that from ReFOLD, were used. Both servers calculate these cavities for a given molecular structure and a defined spherical probe and report their geometrical properties, such as volume, boundary area, buried area, and other parameters described on their websites. In proteins, cavities can be predicted mainly at the barrel end built by an alpha secondary structure, a pattern observed in the refined and unrefined molecule.

### 4.4 LPA and OMPT Docking at the LPA3 receptor binding sites

The 3D configurations of the ligands studied were obtained from PubChem (ID 5497152 for LPA and ID 16078994 for OMPT), and their molecular structure minimization was carried out using the Avogadro program (https://avogadro.cc/docs/tools/auto-optimize-tool/); which allows the net inter-atomic force on each atom to be acceptably close to zero and the position on the potential energy surface is such that the improved structures correspond to the configurations found in nature. The lengths and angles of the links and junctions and their geometry were appropriated and transmitted with optimized energy.

After ligand structure minimization, a blind docking study of the native protein (unrefined) was performed, allowing us to explore the whole molecule. It is worth noticing that it has been reported that Tyr293 participates in the binding of the LPA antagonist, Kil6425, as well as of an extensive series of pharmacological agents used in therapeutics [22]. Residue Trp102 was also particularly interesting because it was consistently found in the hot spot predictions and visualized as exposed and centered in a cavity according to 3D structural analysis with PyMol (https://www.pymol.org/). Therefore, focused docking simulations on the alpha carbon of Tyr293 and Trp102 were also performed. Subsequently, the files of ligands and protein employed to perform the docking analysis were obtained using the program Autodock tools 1.5.6 (ADT) (https://autodocksuite.scripps.edu/adt/). Autogrid4 and Autodock4 programs procured the grid maps and docking results. The PyMol program was also employed to explore the possible ligand-binding cavities in the LPA3 receptor.

Blind docking simulations consist of adjusting the grid box (search area) to the maximum expanse the program allows, i.e., 126 Å3 and a grid space of 0.375 Å3, centered by default on the proteins. All simulations were performed using the LINUX operating system (Centus) and the Lamarckian genetic algorithm, which was chosen to search for the best ligand configurations coupled to the LPA3 receptor. This process generates the most probable results of the ligand conformations within the receptor search area, with a maximum number of 1 x 107 energy evaluations and a population of 100 randomized individuals (represented by the 100 best evaluations). For the focused docking, the grid box was 60 Å3 with a grid space of 0.375 Å3 centered on the alpha carbons of Tyr293 or Trp102, as mentioned previously. The 2D maps showing the receptor’s residues interacting with the ligands were obtained using the Discovery Studio Visualizer (https://discover.3ds.com/discovery-studio-visualizer-download).

### 4.5 Reagents, plasmids, and cells

1-Oleyl lysophosphatidic acid (LPA) and 2S-(1-oleoyl-2-O-methyl-glycerophosphothionate) (OMPT) were from Cayman Chemical Co. (Ann Arbor, MI, USA). Dulbecco’s modified Eagle’s medium, trypsin, Lipofectamine 2000, streptomycin, penicillin, amphotericin B, blasticidin, hygromycin B, doxycycline hyclate, and Fura-2 AM were purchased from Invitrogen-Life Technologies (Carlsbad, CA, USA). Our previous publications indicate the source of other materials [28, 32, 33]. Parental Flp-In T-Rex HEK293 cells were obtained from Invitrogen (Carlsbad, CA, USA). The LPA3 receptor sequence was fused at the carboxyl terminus (Ctail) with a green fluorescent protein and cloned into the pCDNA5/FRT/TO plasmid to employ the inducible Flp-In TREx expression system [28, 32, 33]. Mutants of this plasmid were obtained commercially (Bioinnovatise, Inc., Rockville, MD, USA), and the changes were confirmed by sequencing. Those amino acids identified to interact with the ligands in the docking simulations were substituted by alanine. Two main agonist-binding cavities were detected (Cavity 1 and 2, see Results). The residues replaced by alanine for the Upper Cavity mutant were (Thr25, Leu86, Thr90, Arg105, Gln106, Asp110, Leu113, Leu179, Trp191, Trp252, Gly255, Leu259, Lys275, Arg276, Phe278, and Leu279). For the Lower Cavity mutant, the amino acids substituted were (Ile131, Met134, Arg135, Val136, Val213, Lys216, Thr217, Val219, Leu220, Arg230, Pro234, Lys275, Arg276, and Phe278). Only the Lower Cavity’s last three amino acids (i.e., Lys275, Arg276, and Phe278) were present in the Upper Cavity. A third mutant was also obtained in which all the residues detected in these binding pockets were substituted by alanine (Upper+Lower Cavity mutant). T-Rex HEK293 cells were transfected with plasmids for the expression of the wild-type and mutant receptors and were subjected to selection as described before [28, 32, 33].

### 4.6 Intracellular Calcium Concentration

Determinations were performed as previously described [28, 32, 33]. In brief, the cells were serum-starved and treated for 12 h with 100 ng/mL doxycycline hyclate to induce LPA3 expression; all receptors (wild-type and mutants were tagged with the green fluorescent protein [28, 32, 33]; the density was similar as reflected by fluorescence microscopy). The cells were loaded with 2.5 μM Fura-2 AM for 1 h. Cells were carefully detached from the Petri dishes, washed to eliminate unincorporated dye, and maintained in suspension. Two excitation wavelengths (340 and 380 nm) and an emission wavelength of 510 nm were employed. Intracellular calcium levels were calculated as described by Grynkiewicz et al.[34].

### 4.7 Statistical Analyses

The data are presented as the means ± standard error of the means. Statistical analyses were performed using ordinary one-way ANOVA with the Bonferroni post-test using the software included in the GraphPad Prism program (version 10.2.3). A p-value < 0.05 was considered statistically significant.

## Supplementary Materials

The following supporting information can be downloaded at: www.mdpi.com/xxx/s1, Figure S1: title; Table S1: title; Video S1: title.

## Author Contributions

Conceptualization: K.H.S., J.C.-B., J.A.G.-S.; Methodology: K.H.S., J.C.-B.; Investigation: K.H.S., M.T.R.-A., R.R.-H., J.C.-B.; Visualization: K.H.S., M.T.R.-A., R.R.-H., J.C.-B.; Writing – original draft: K.H.S., J.C.-B., J.A.G.-S.; Writing – review & editing: K.H.S., M.T.R.-A., R.R.-H., J.C.-B., J.A.G.-S.

## Data Availability Statement

The data are available from the corresponding author upon reasonable request.

## Acknowledgments

We thank Dr. A. César Poot-Hernández for his suggestions to improve the manuscript. The advice and technical support of the following members of the indicated Service Units of our Institute is gratefully acknowledged: Dr. Héctor Malagón and Dr. Claudia Rivera (Bioterio); Juan Barbosa and Gerardo Coello (Cómputo); and Aurey Galván and Manuel Ortínez (Taller). K. Helivier Solís is a student of the Programa de Doctorado en Ciencias Biomédicas (UNAM) of Universidad Nacional Autónoma de México (account 520014983).

## SUPPLEMENTARY MATERIAL

**Supplementary Figure S1.**
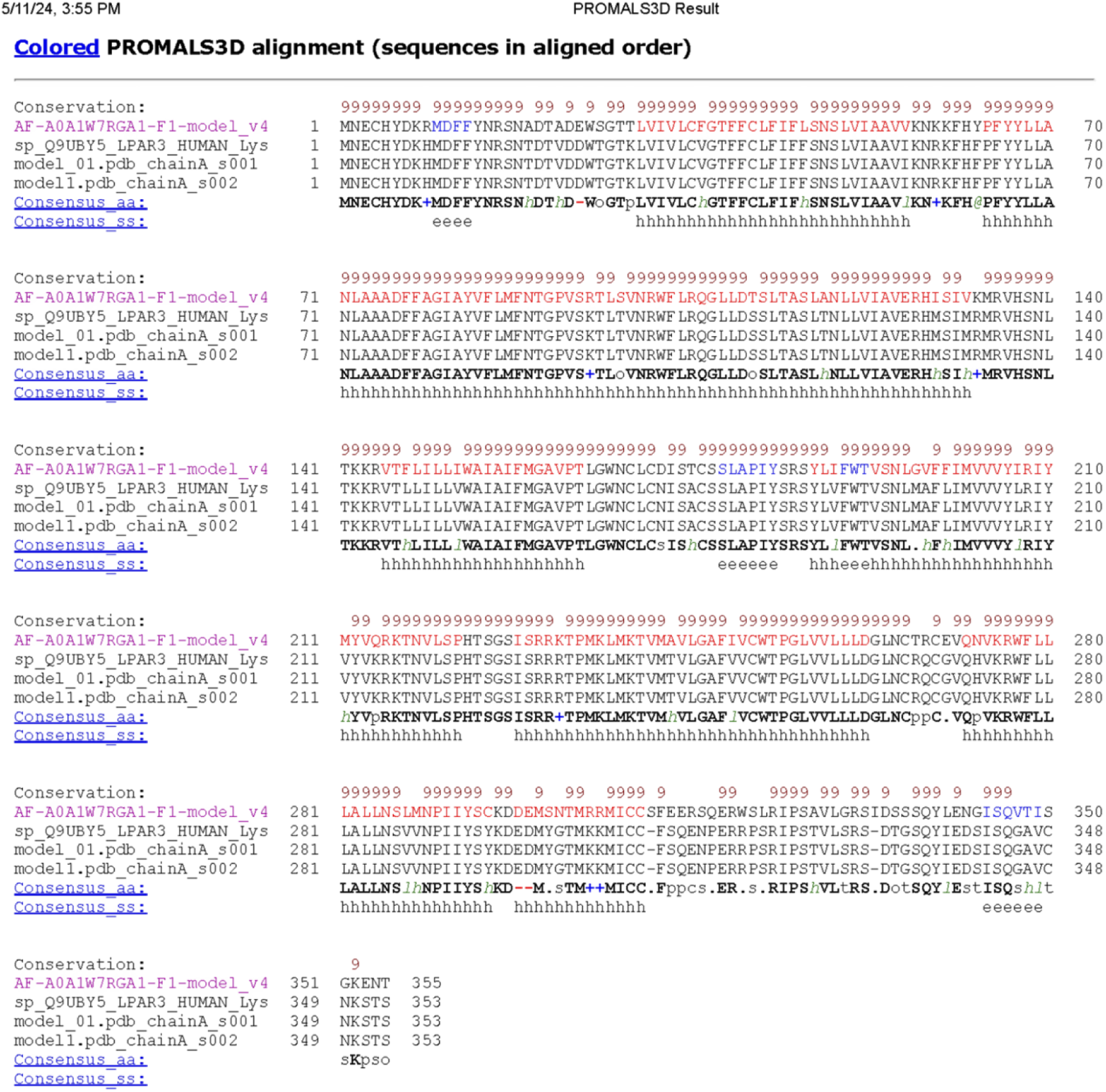
Alignment of the FASTA sequence of the LPA_3_ receptor (Q9UBY5) with the 3D model programs I-Tasser (model 01.pdb chainA_s001), Swiss Model (model1) and AlphaFold (AF-A v4).

**Supplementary Figure S2.**
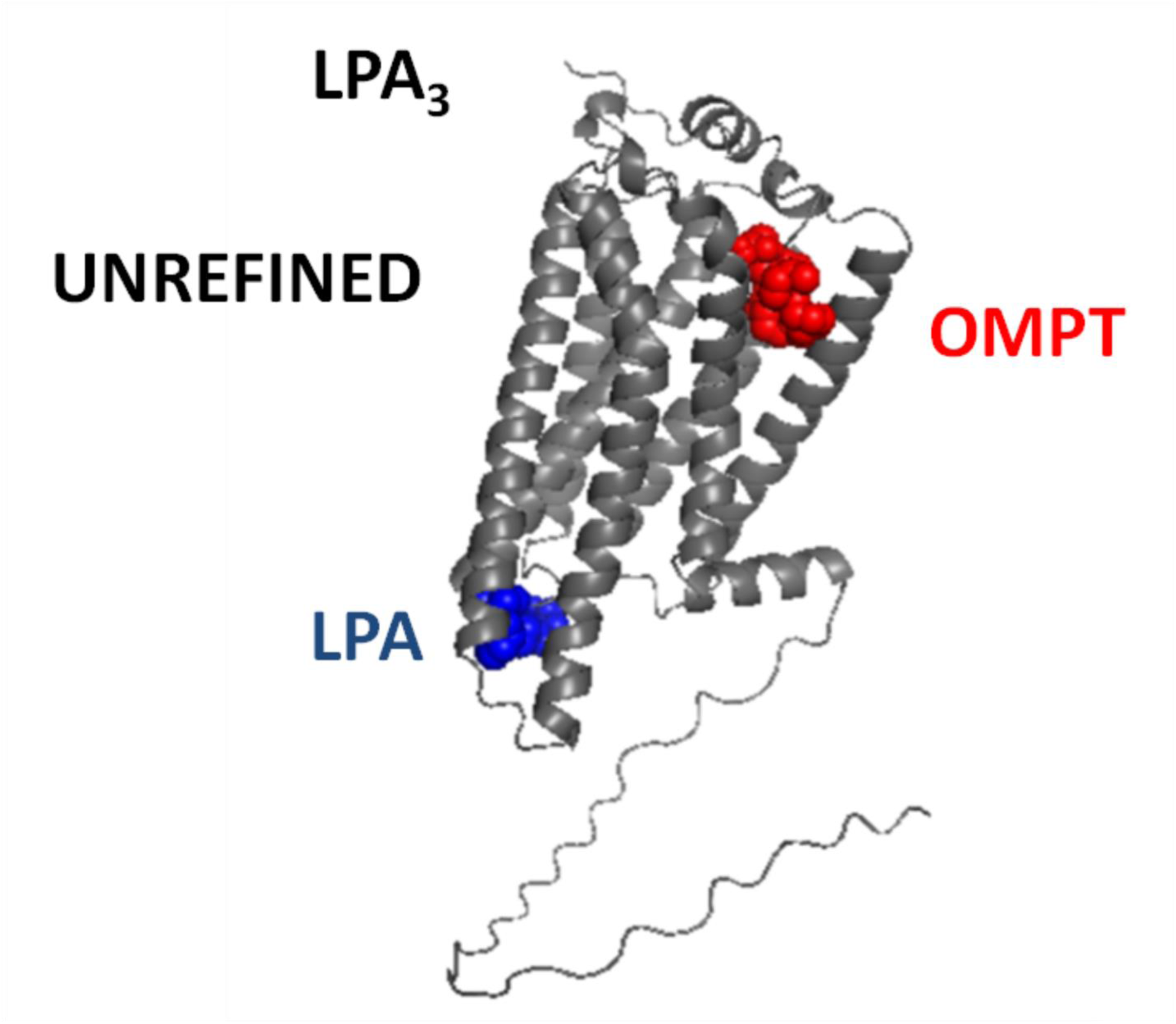
Images resulting from the LPA_3_-LPA (blue) and LPA_3_-OMPT (red) docking analysis using PyMol.

**Supplementary Figure S3.**
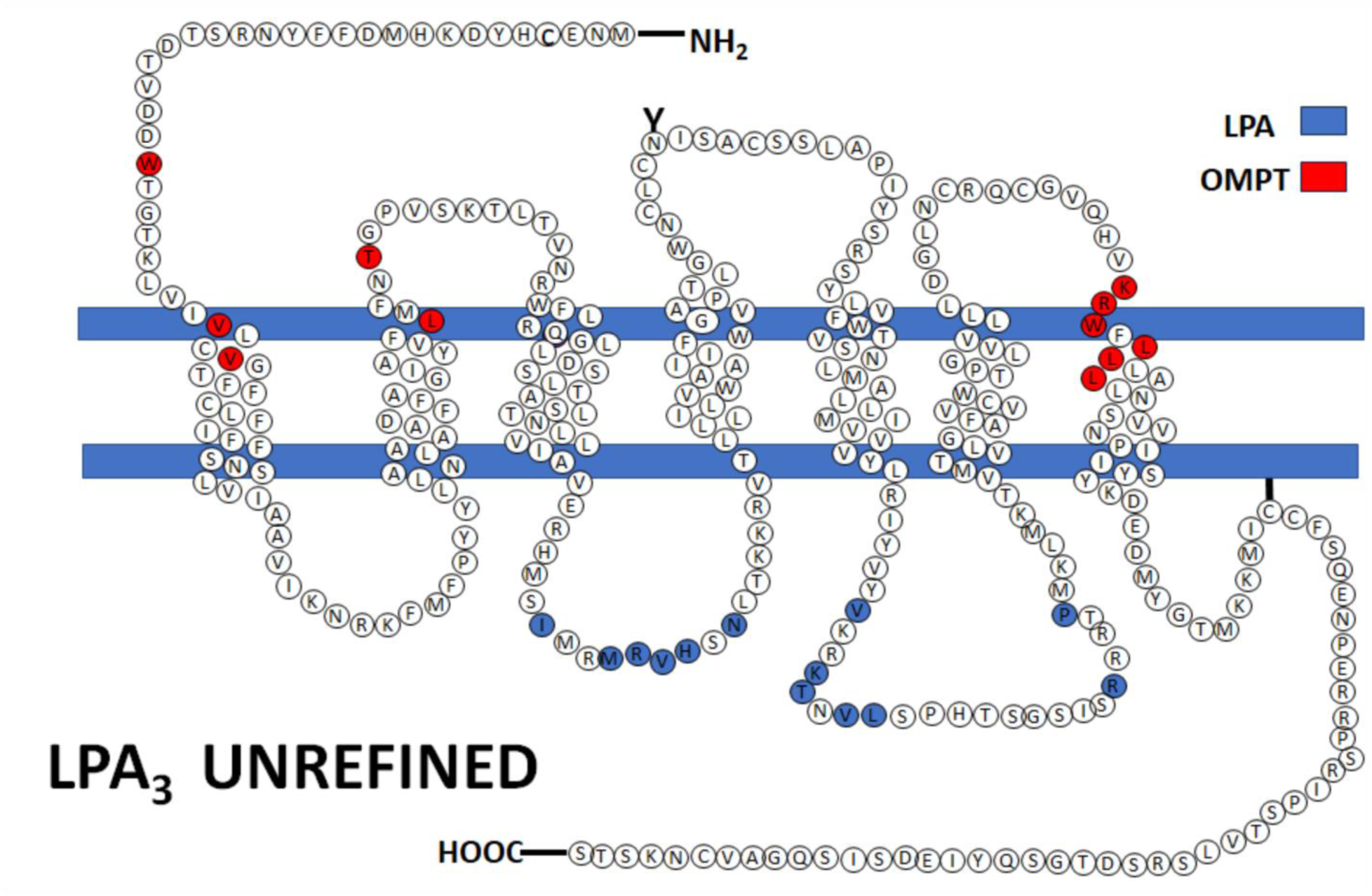
Cartoon showing the LPA_3_ receptor sites interacting with LPA (blue) and OMPT (red).

**Supplementary Figure S4.**
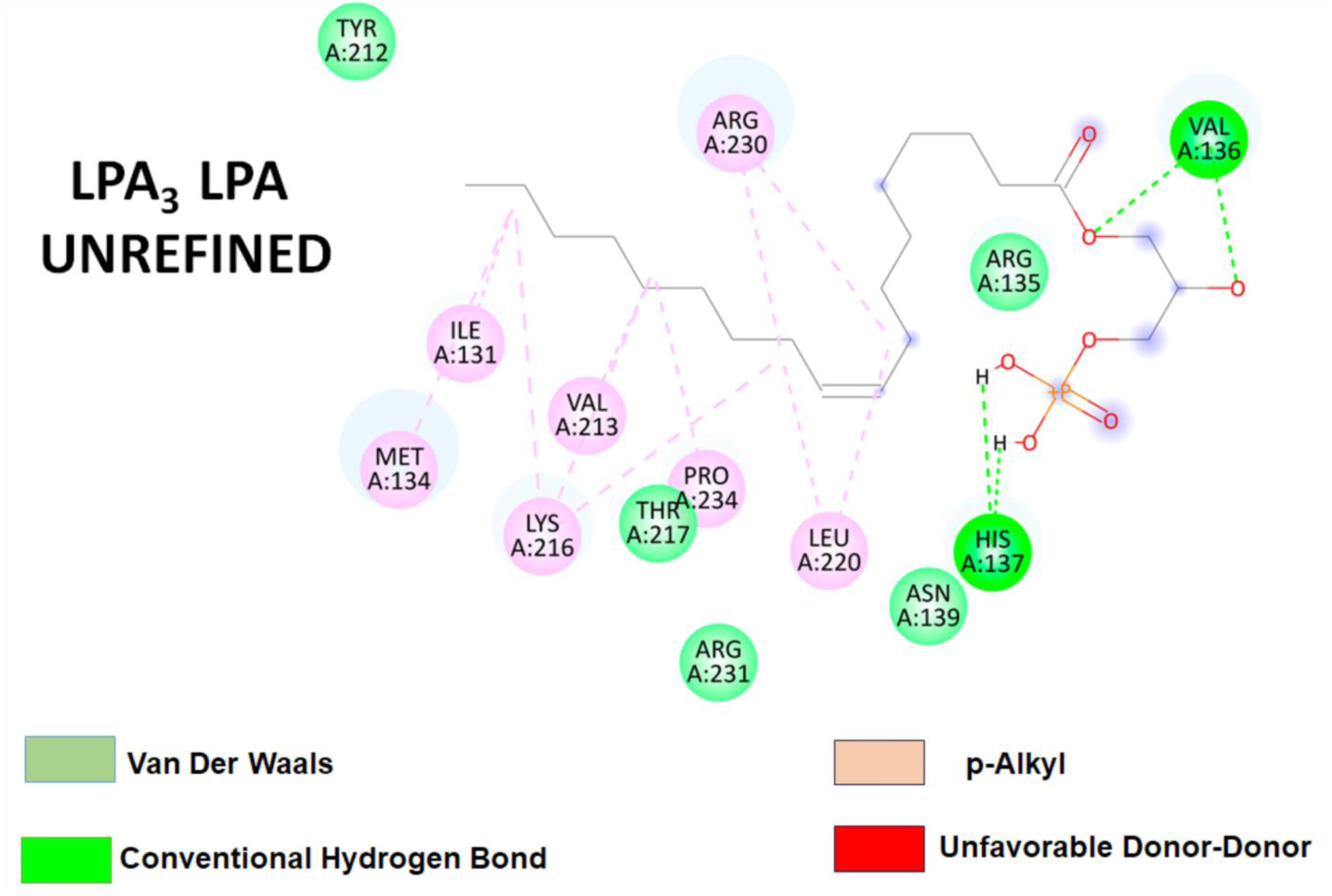
The 2D diagram shows interactions observed at the LPA_3_ binding site (Lower Cavity) with LPA (Discovery Studio Visualizer); critical amino acids contributing to interactions are shown in circles.

**Supplementary Figure S5.**
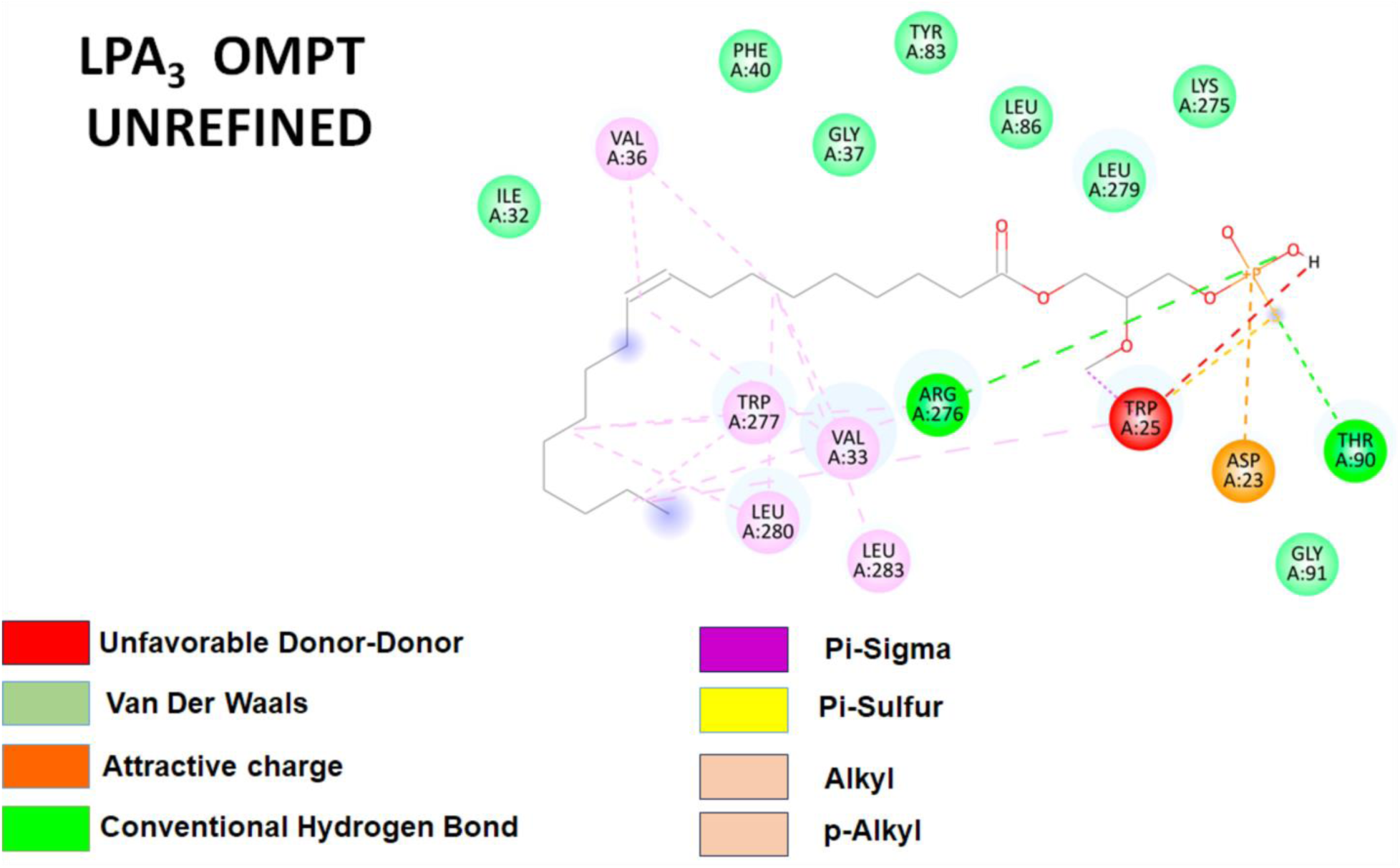
The 2D diagram shows interactions at the LPA_3_ binding site (Upper Cavity) with OMPT (Discovery Studio Visualizer). Critical amino acids contributing to interactions are shown in circles.

**Supplementary Figure S6.**
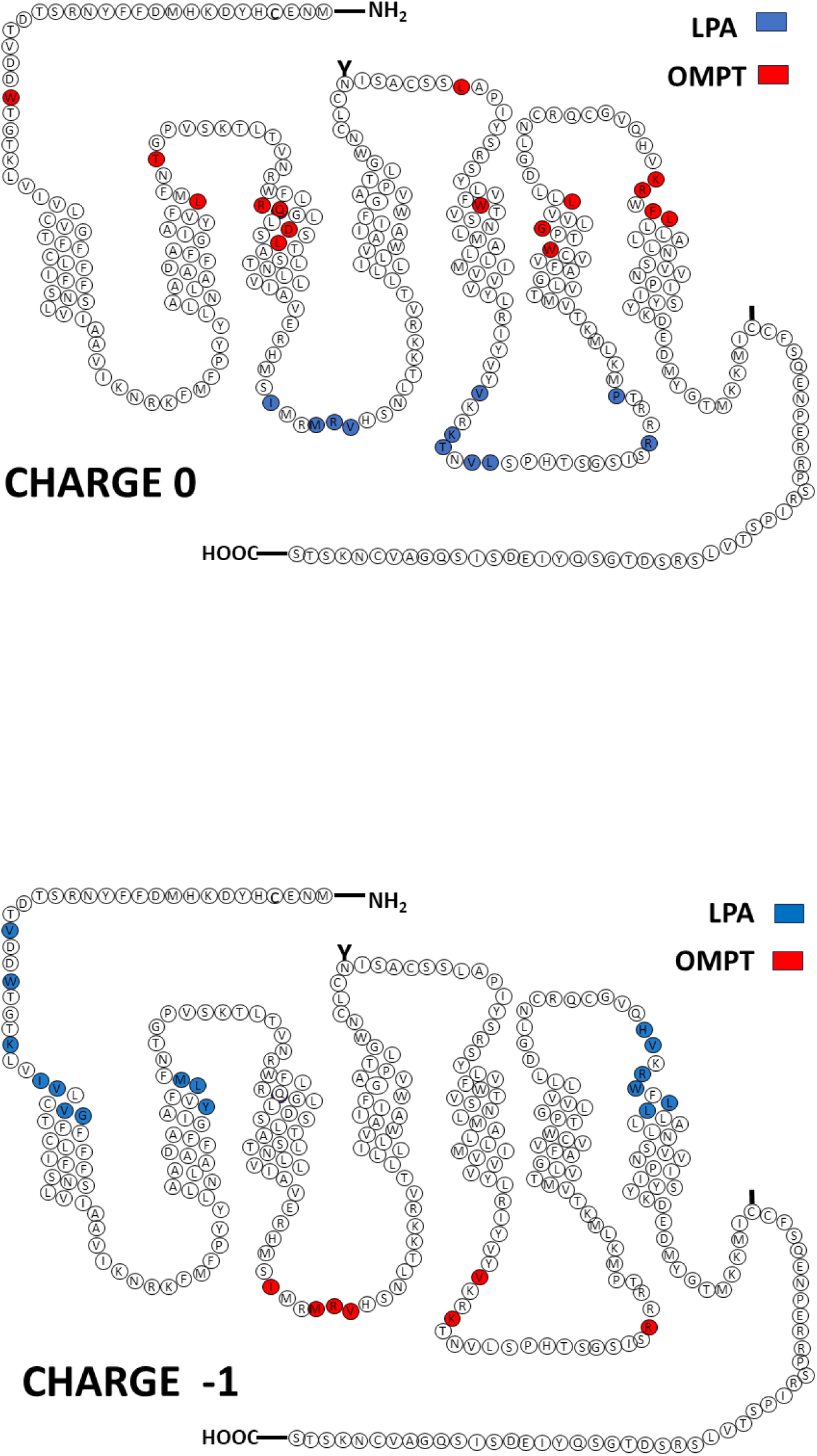
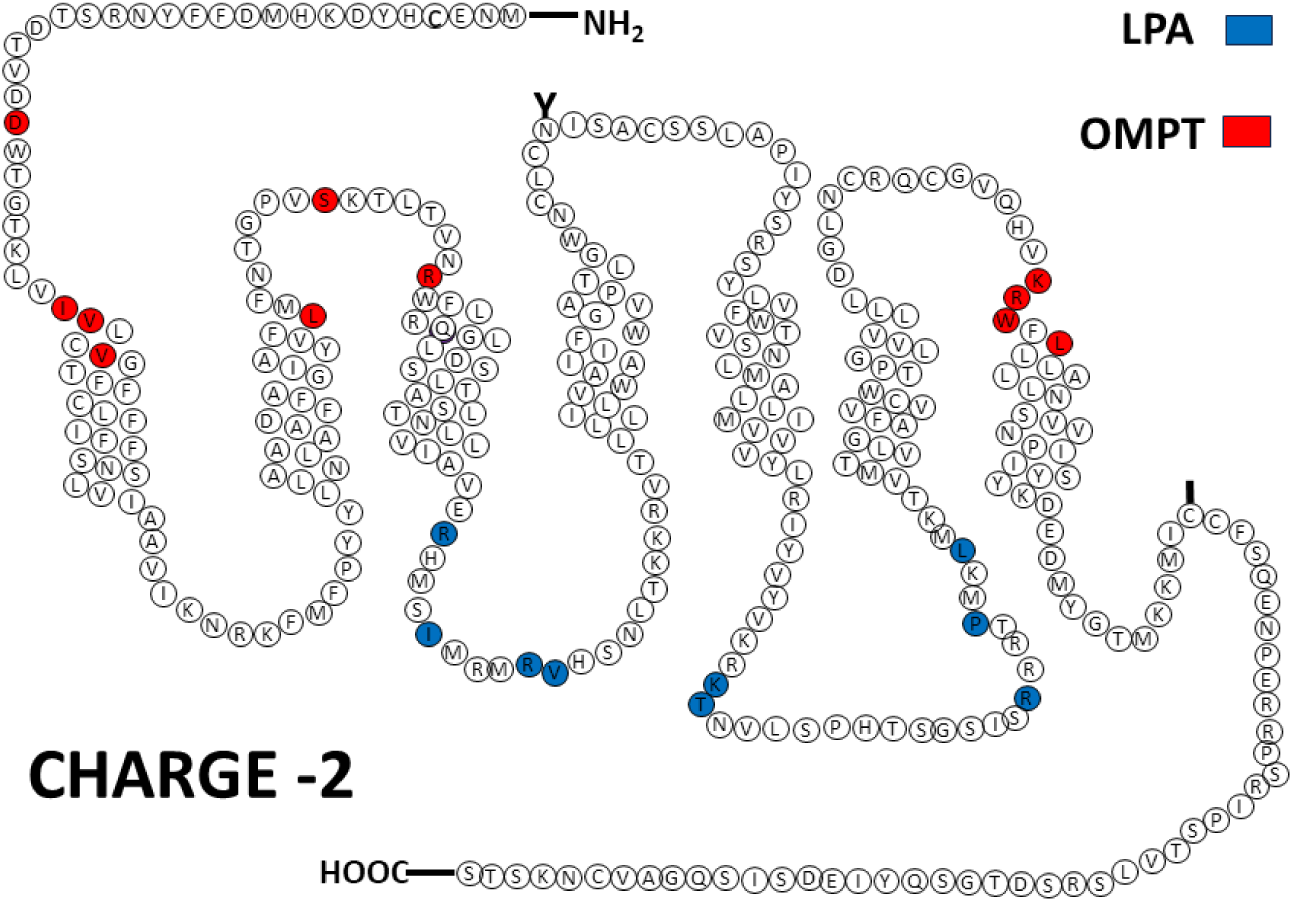
Cartoons showing the refined LPA_3_ receptor sites interacting with LPA (blue) and OMPT (red). Charges are indicated in the Figures

**Supplementary Figure S7.**
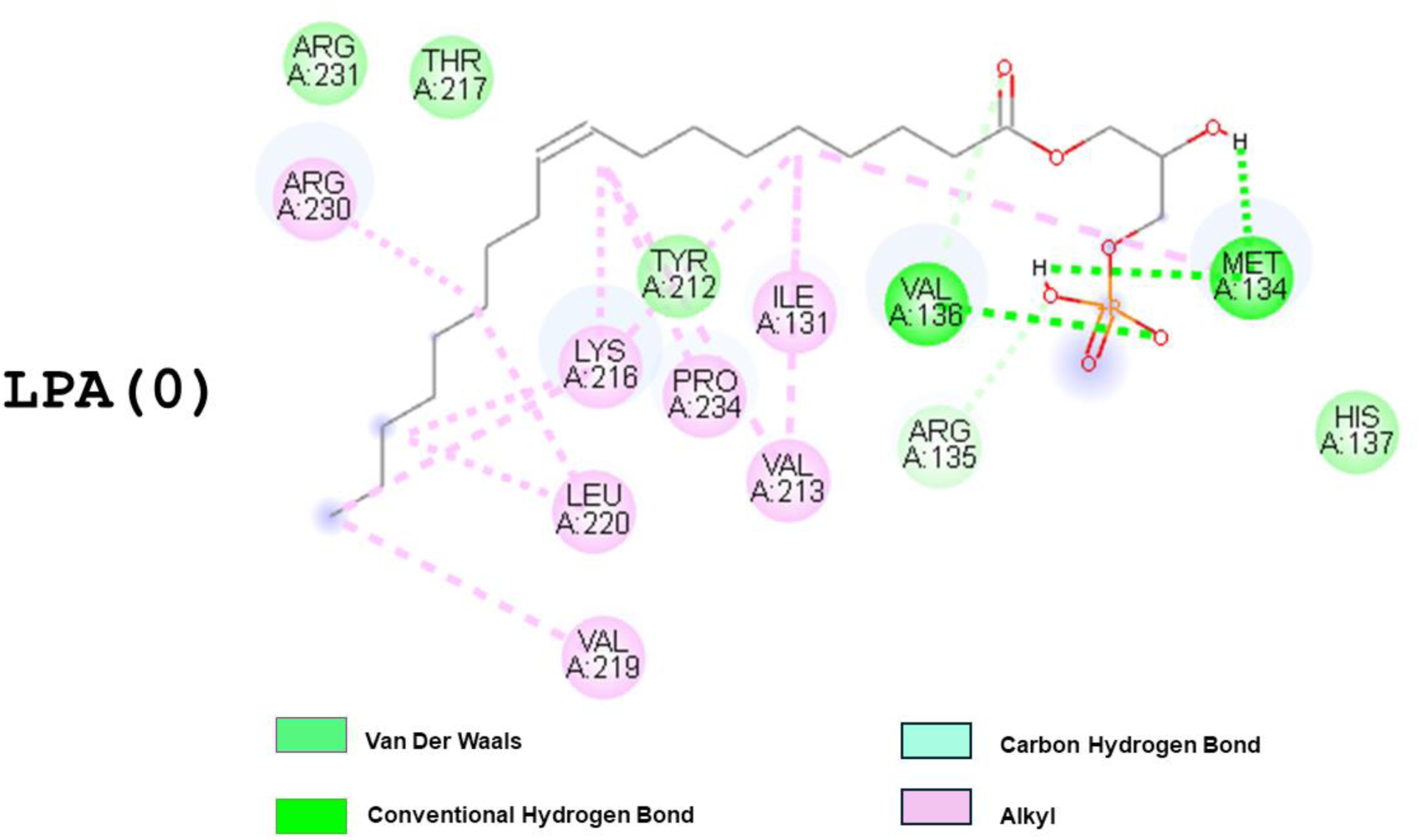
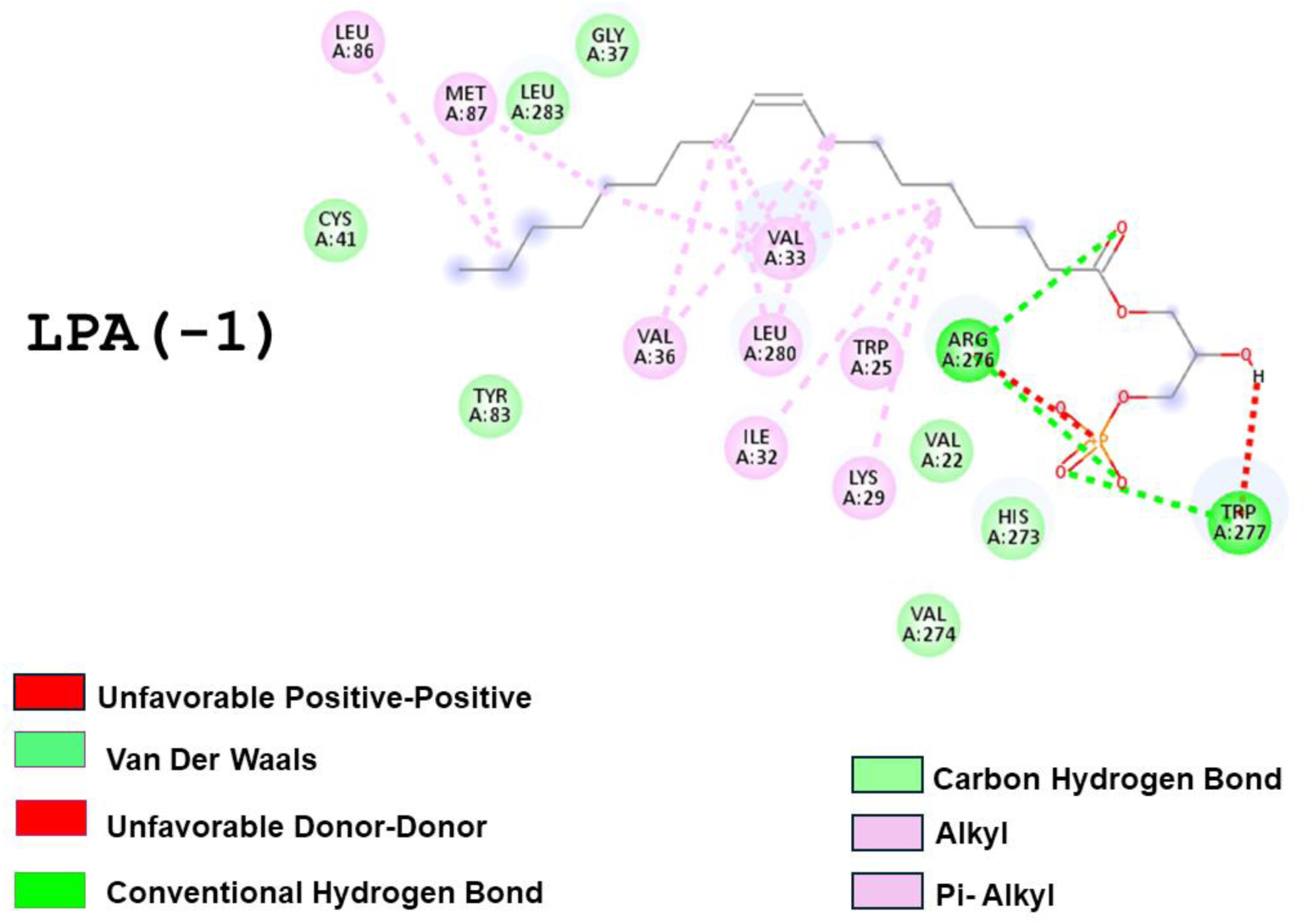
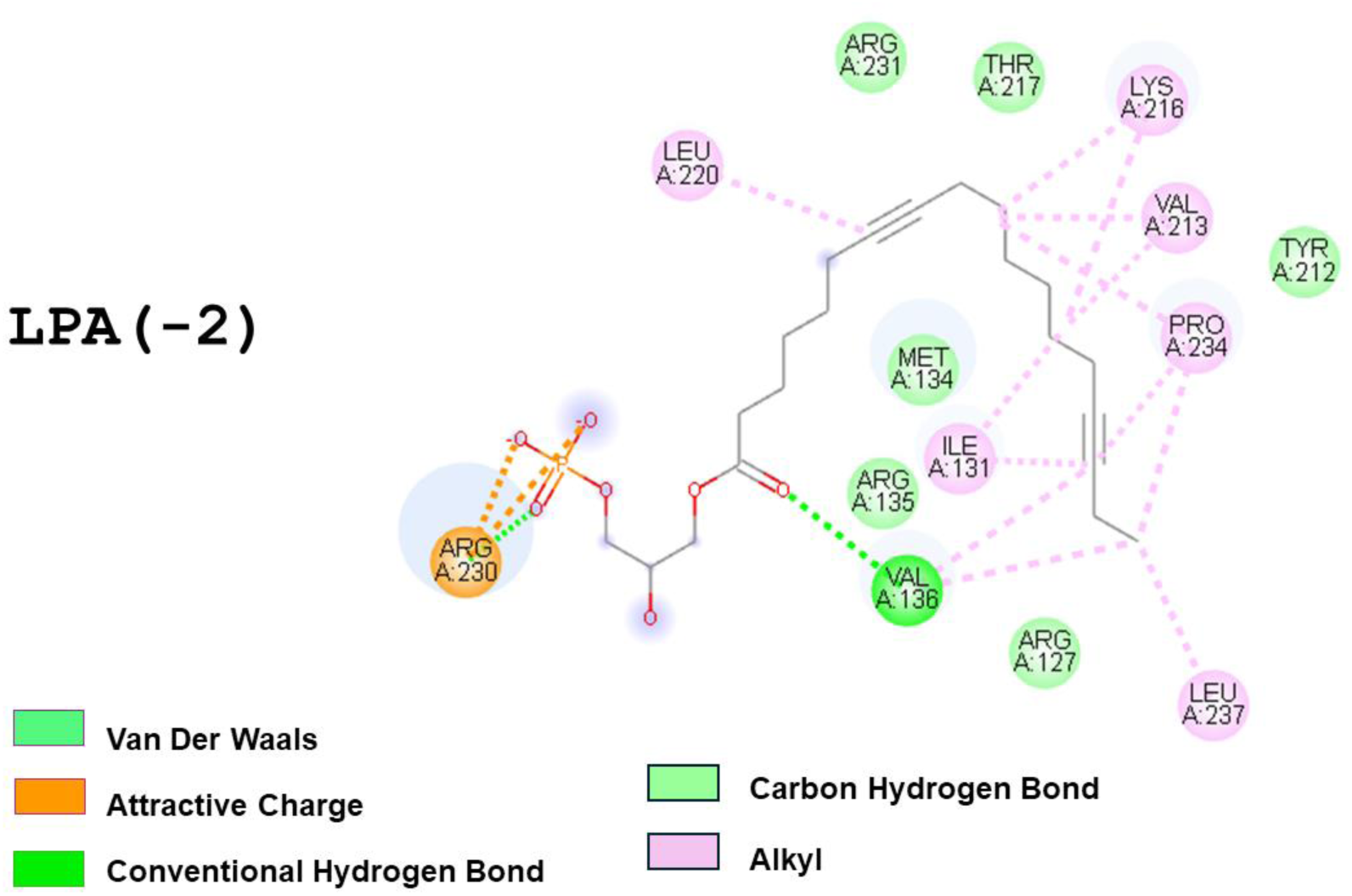
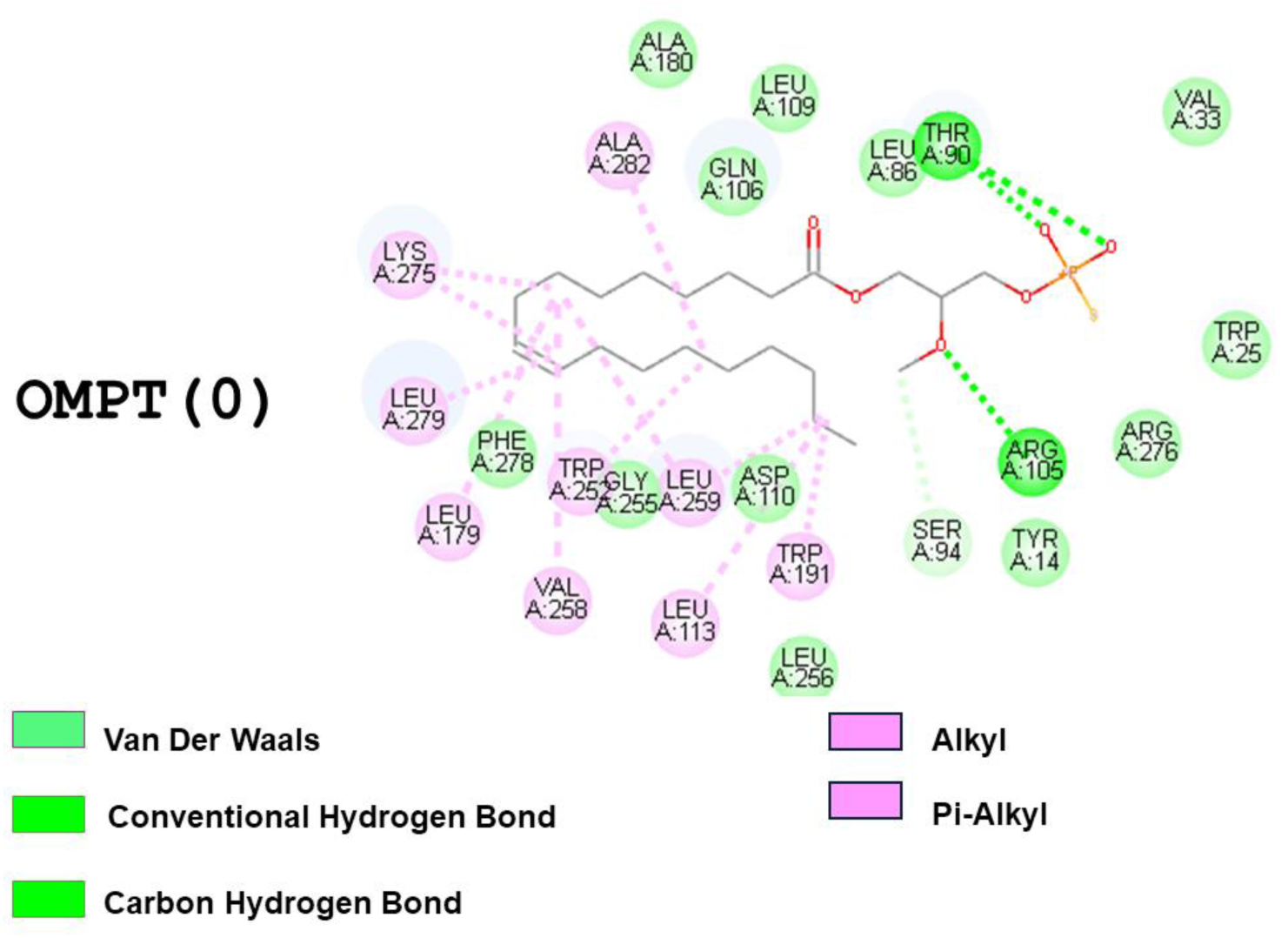
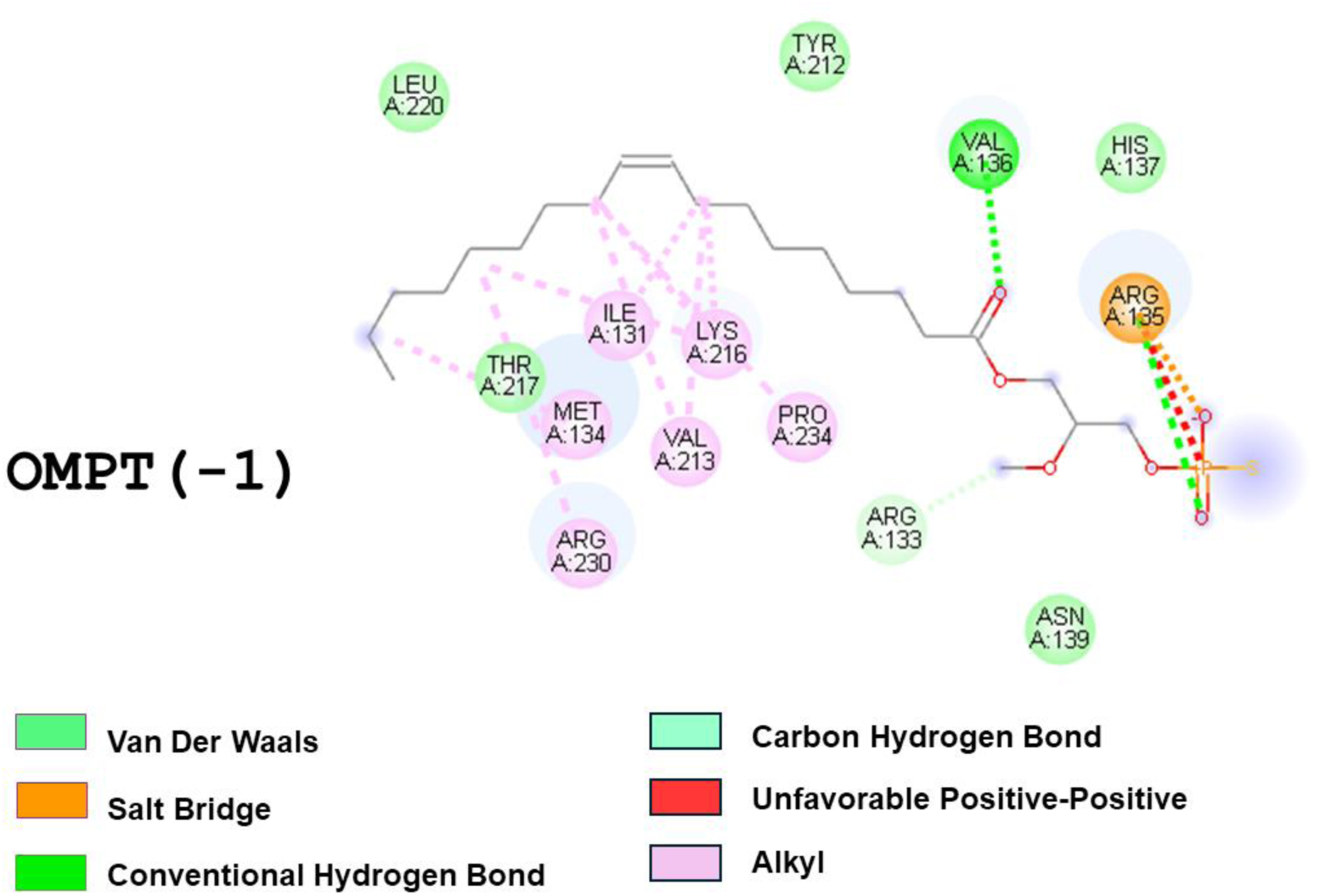
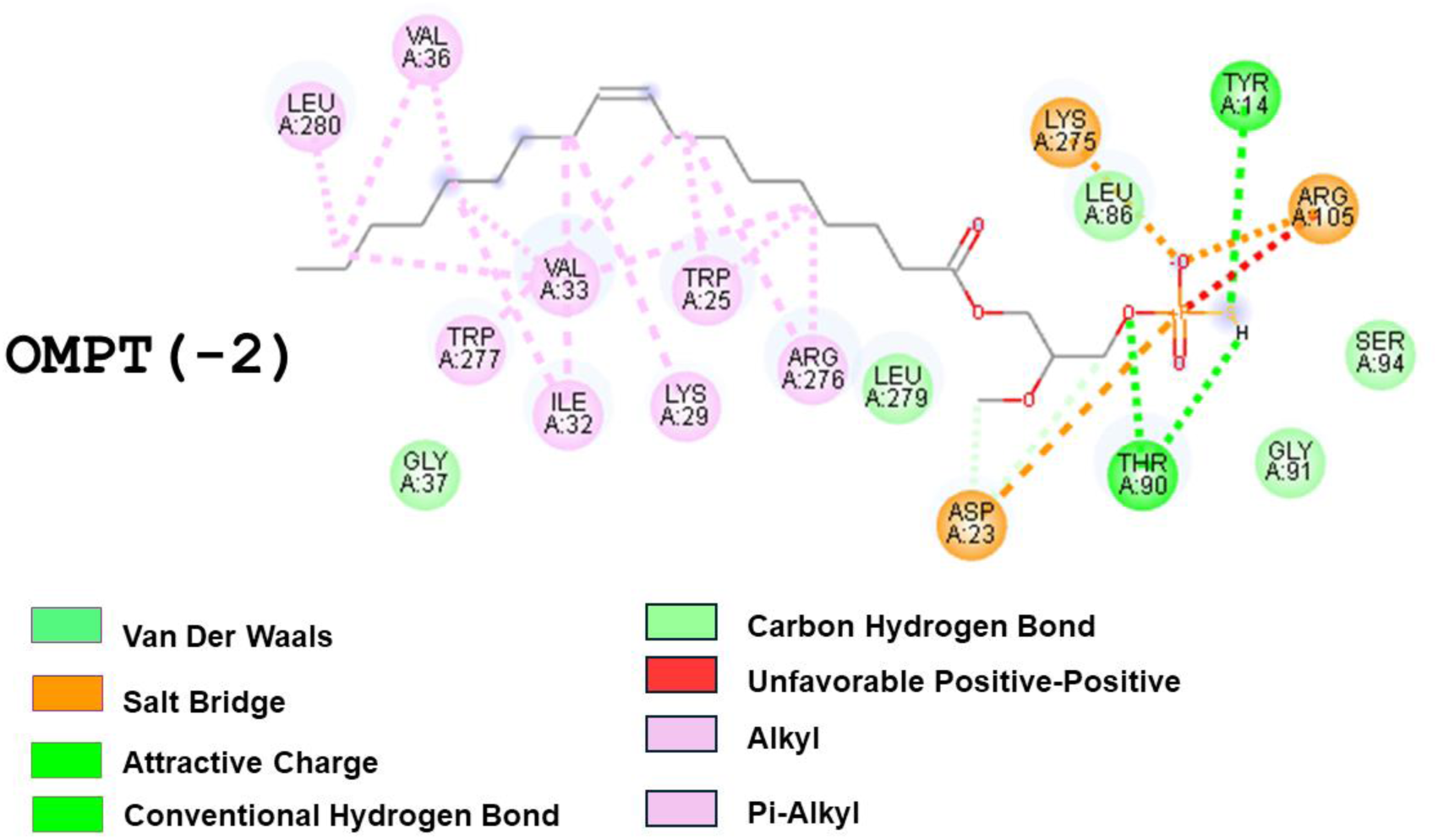
The 2D diagram shows interactions at the LPA_3_ binding site with LPA or OMPT. In the individual images Ligand and Charge are indicated (Discovery Studio Visualizer). Critical amino acids contributing to interactions are shown in circles.

**Supplementary Figure S8.**
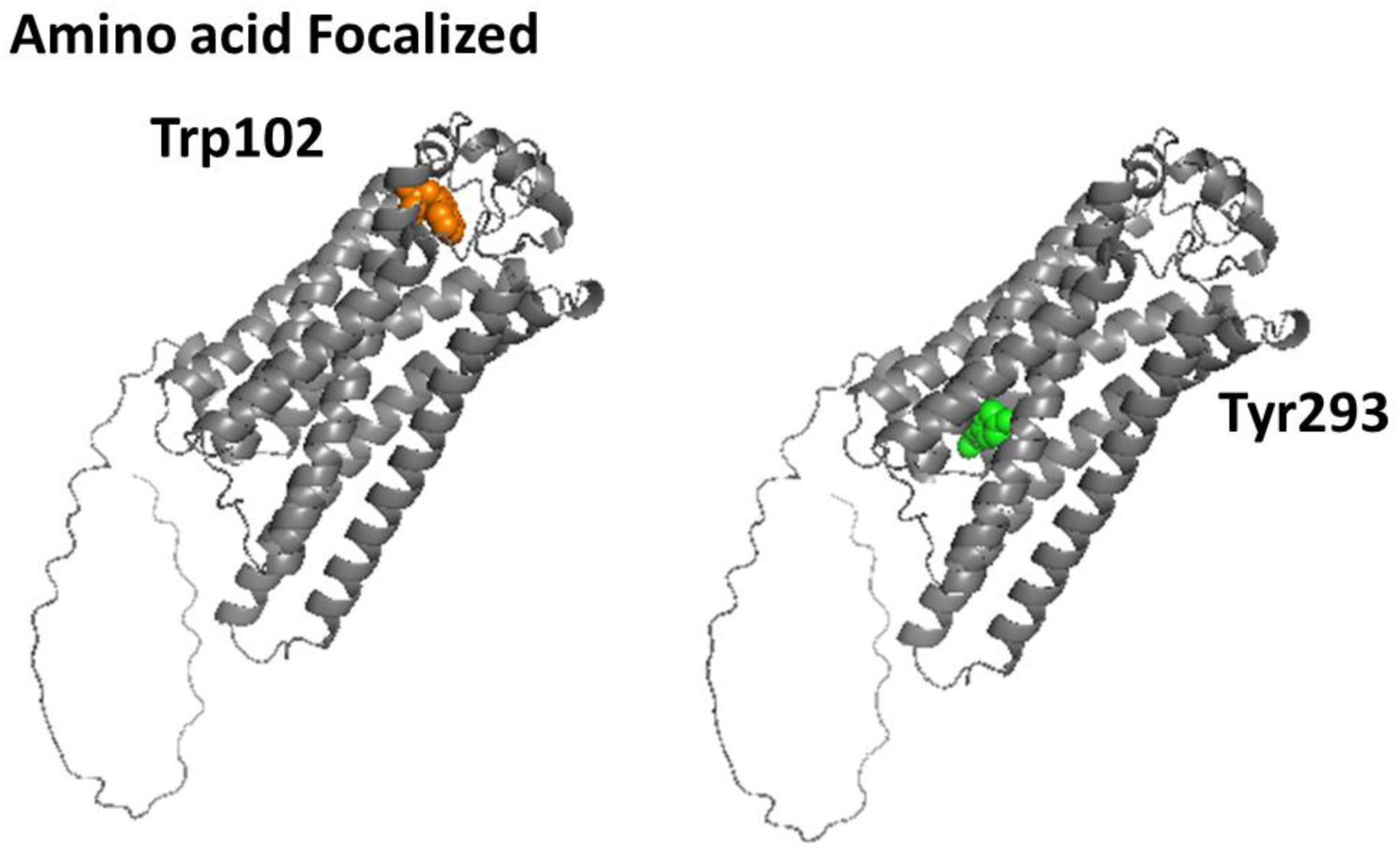
Localization of the amino acids at the LPA_3_ receptor to be focused. Trp102, orange; Tyr293, green.

**Supplementary Figure S9.**
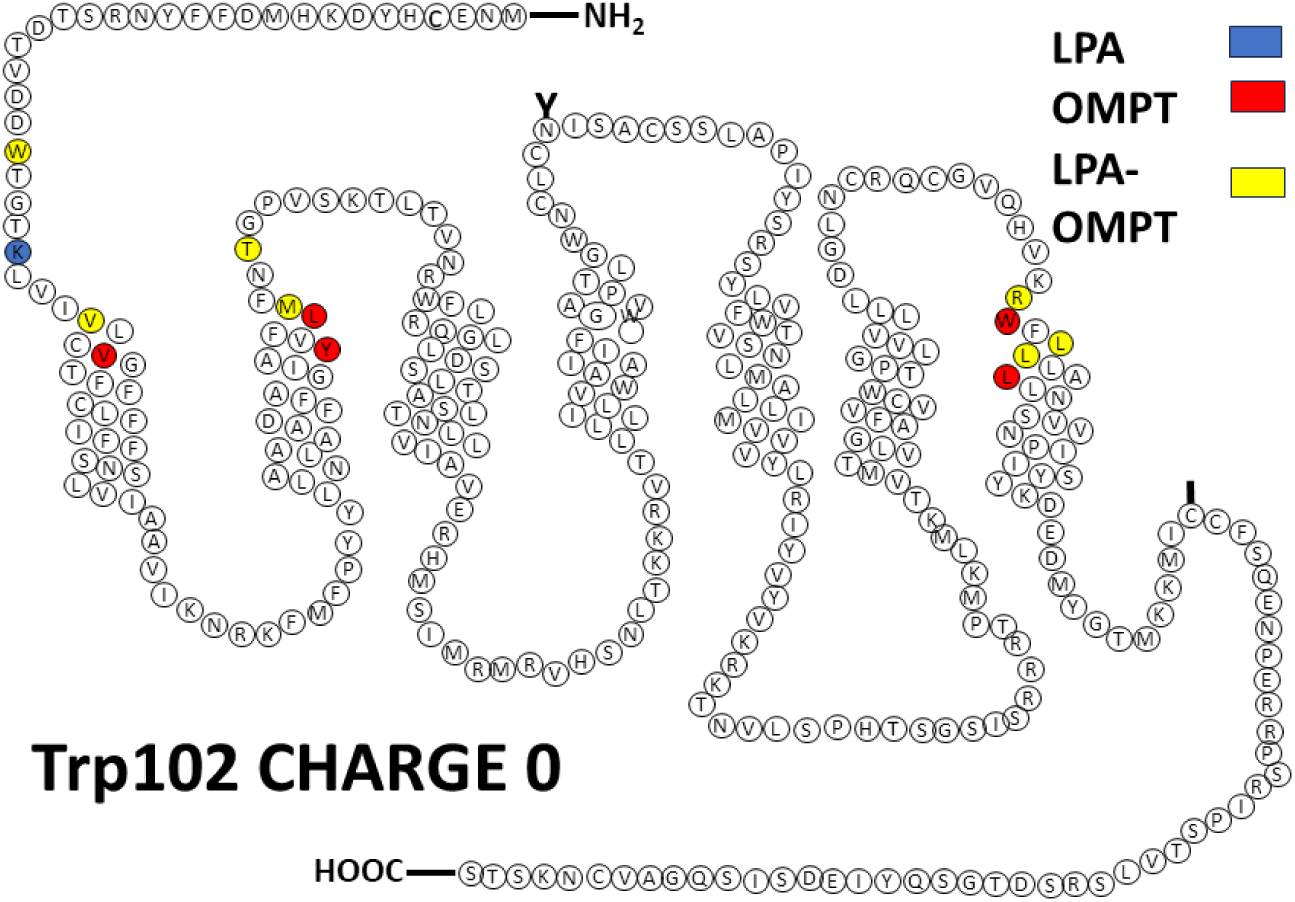
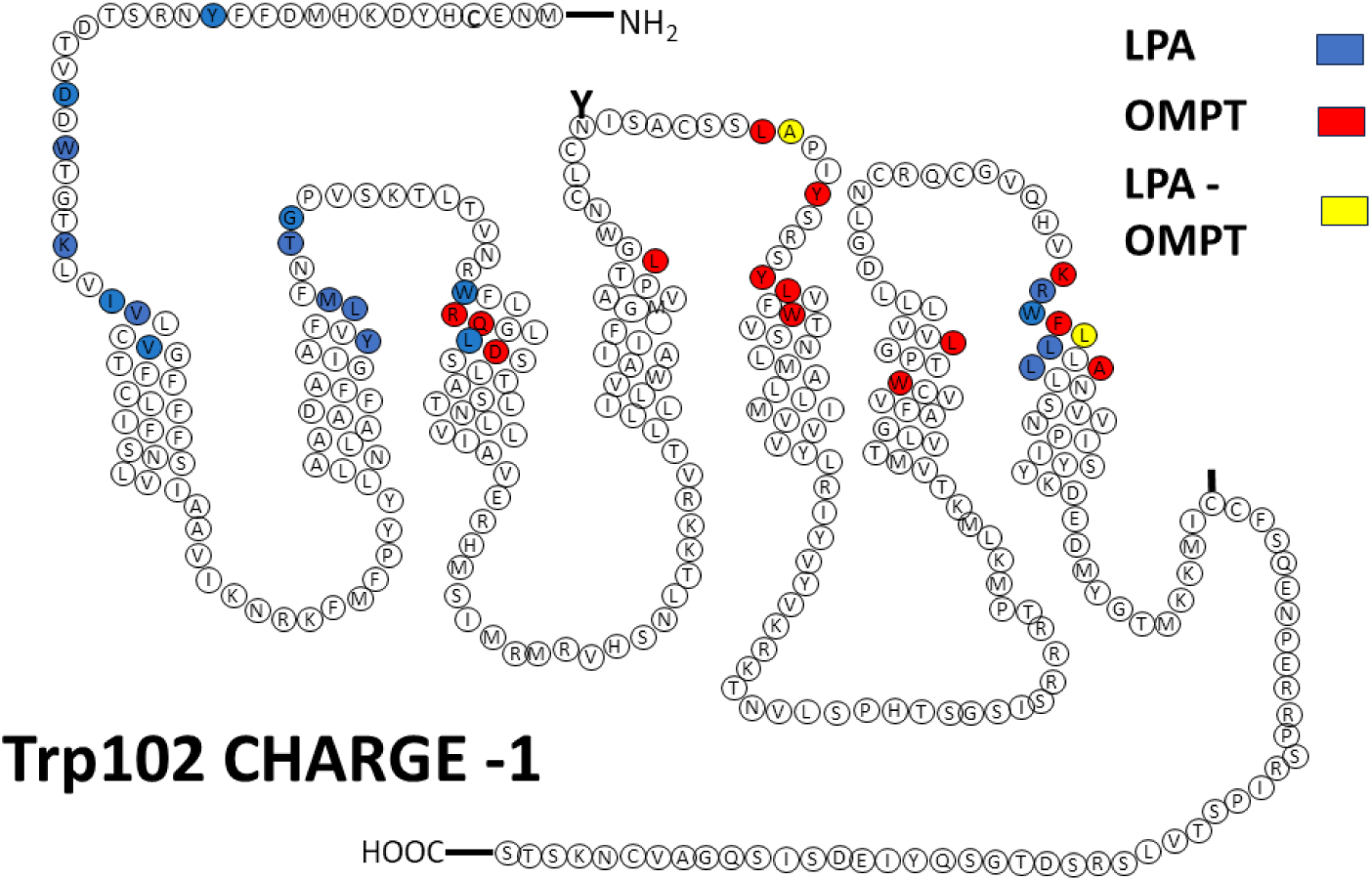
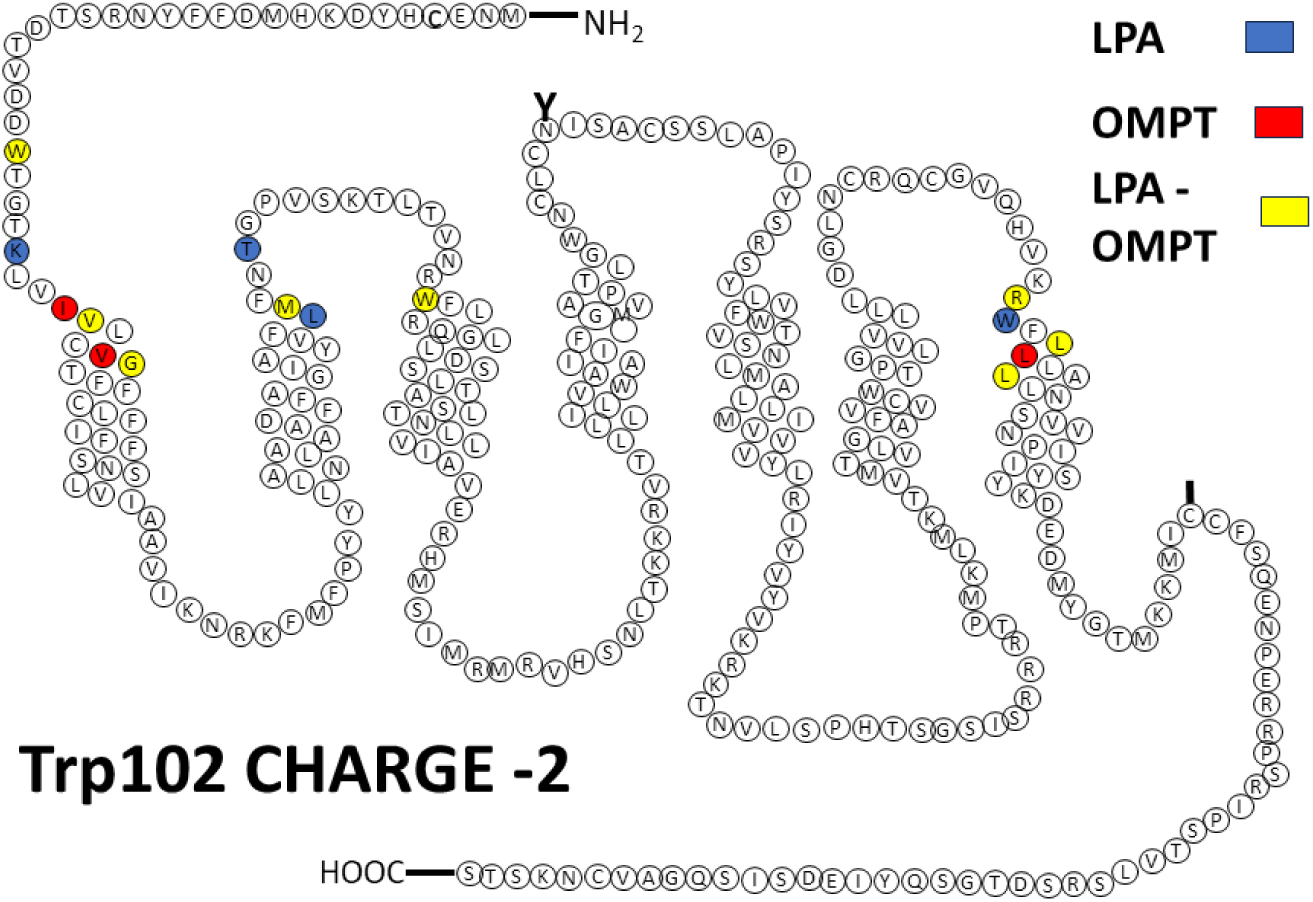
Cartoons showing the LPA_3_ receptor (Trp102-focused) sites interacting with LPA (blue) and OMPT (red).Charges are indicated in the Figures

**Supplementary Figure S10.**
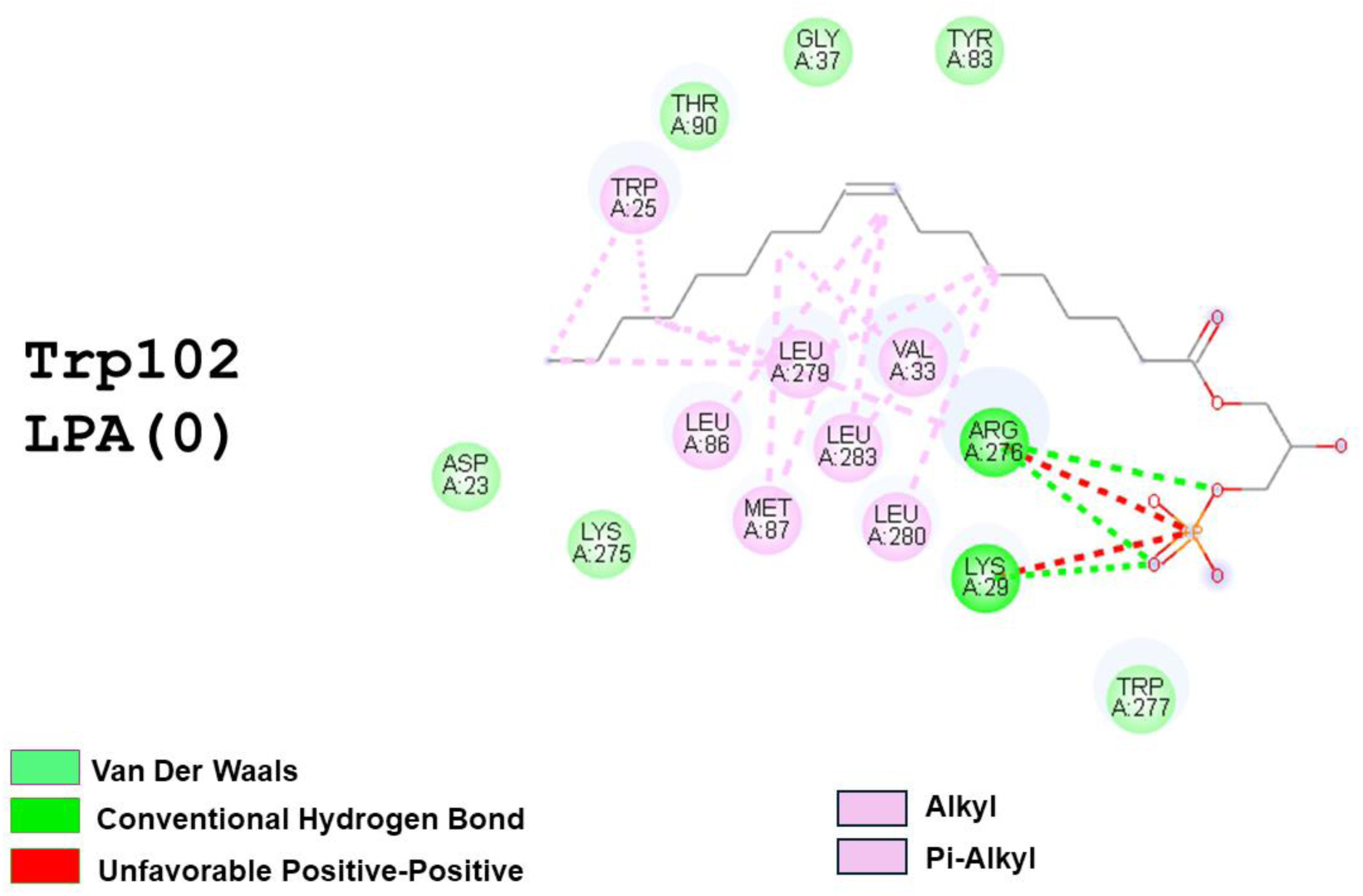
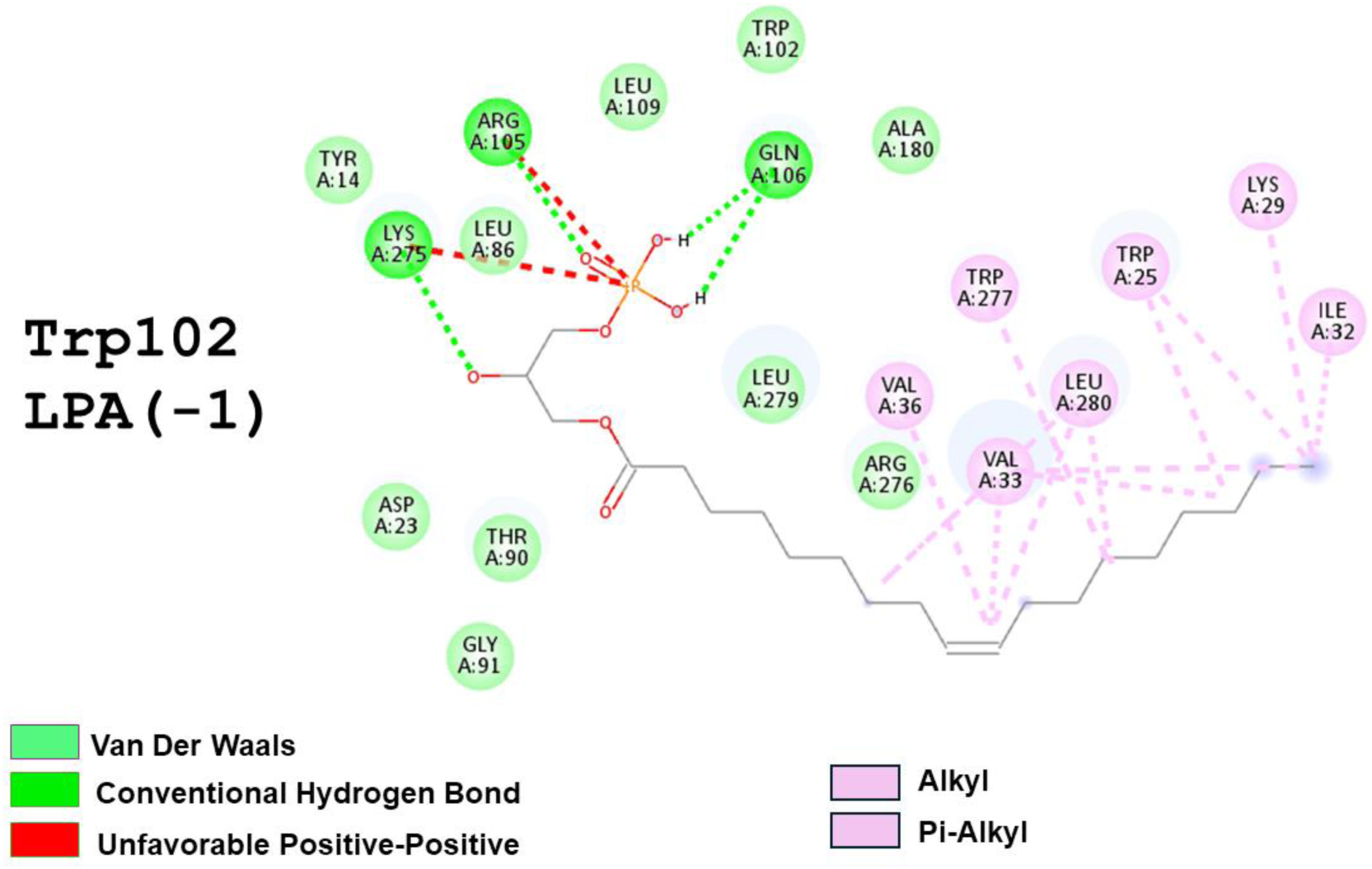
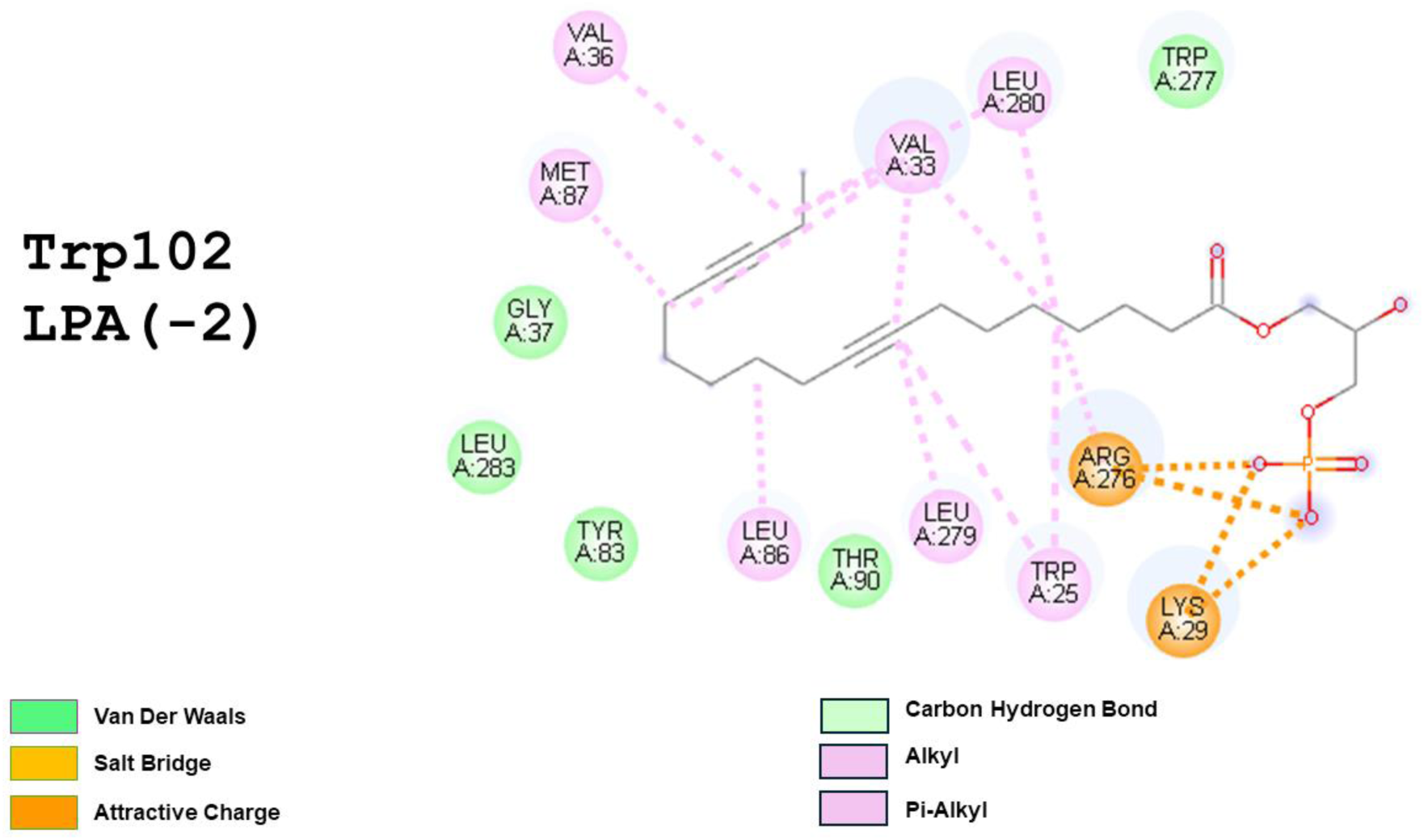
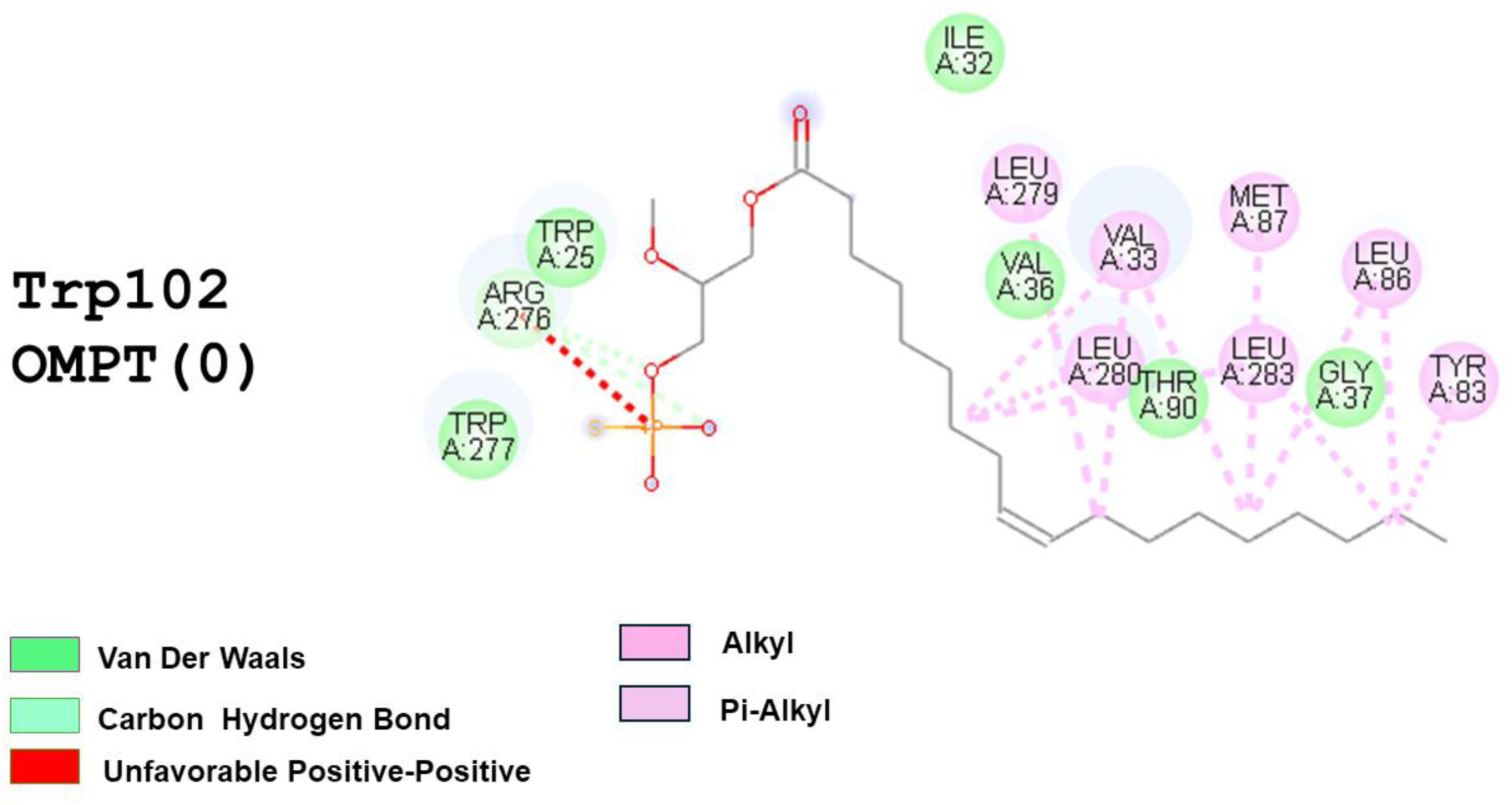
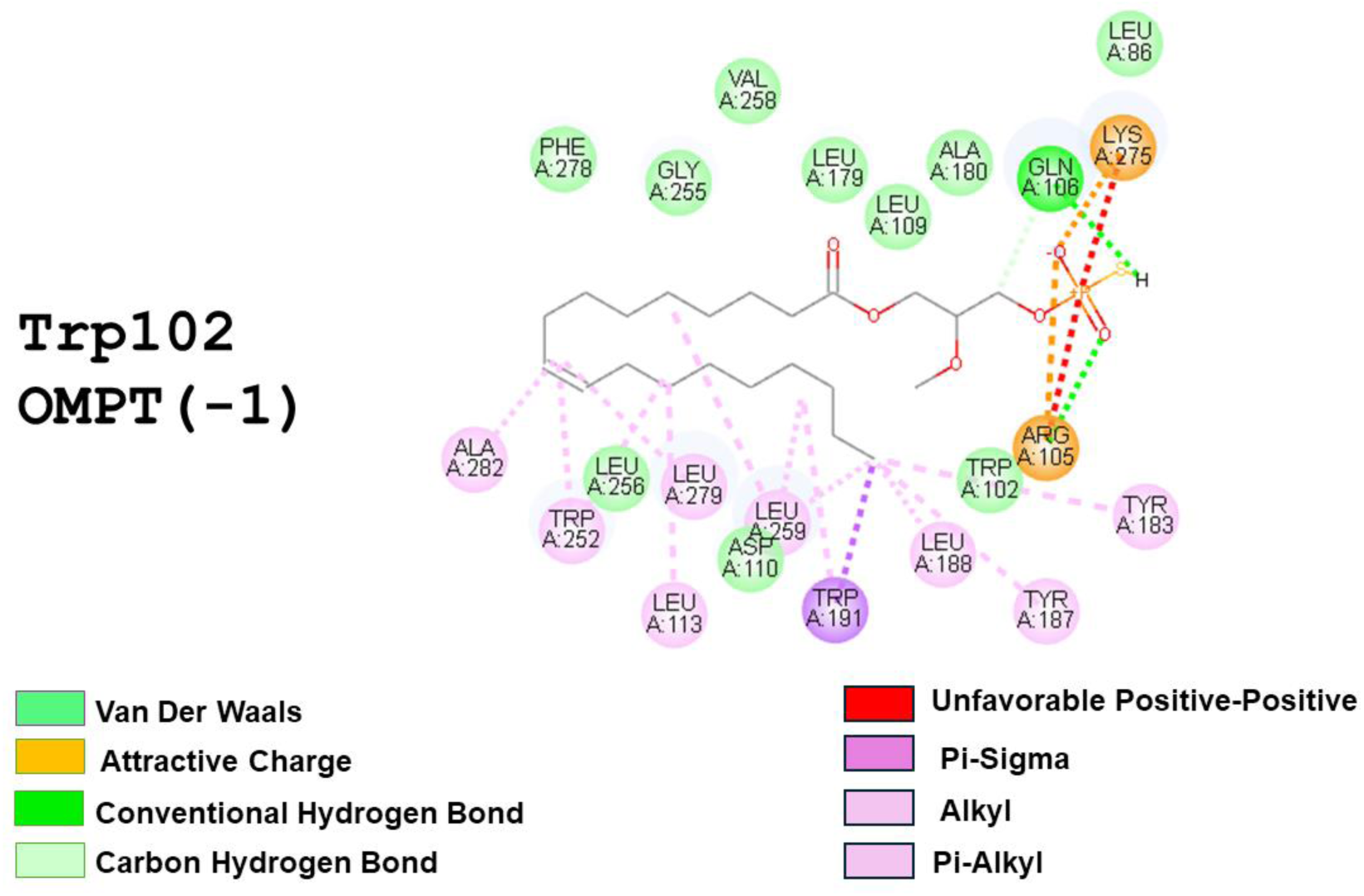
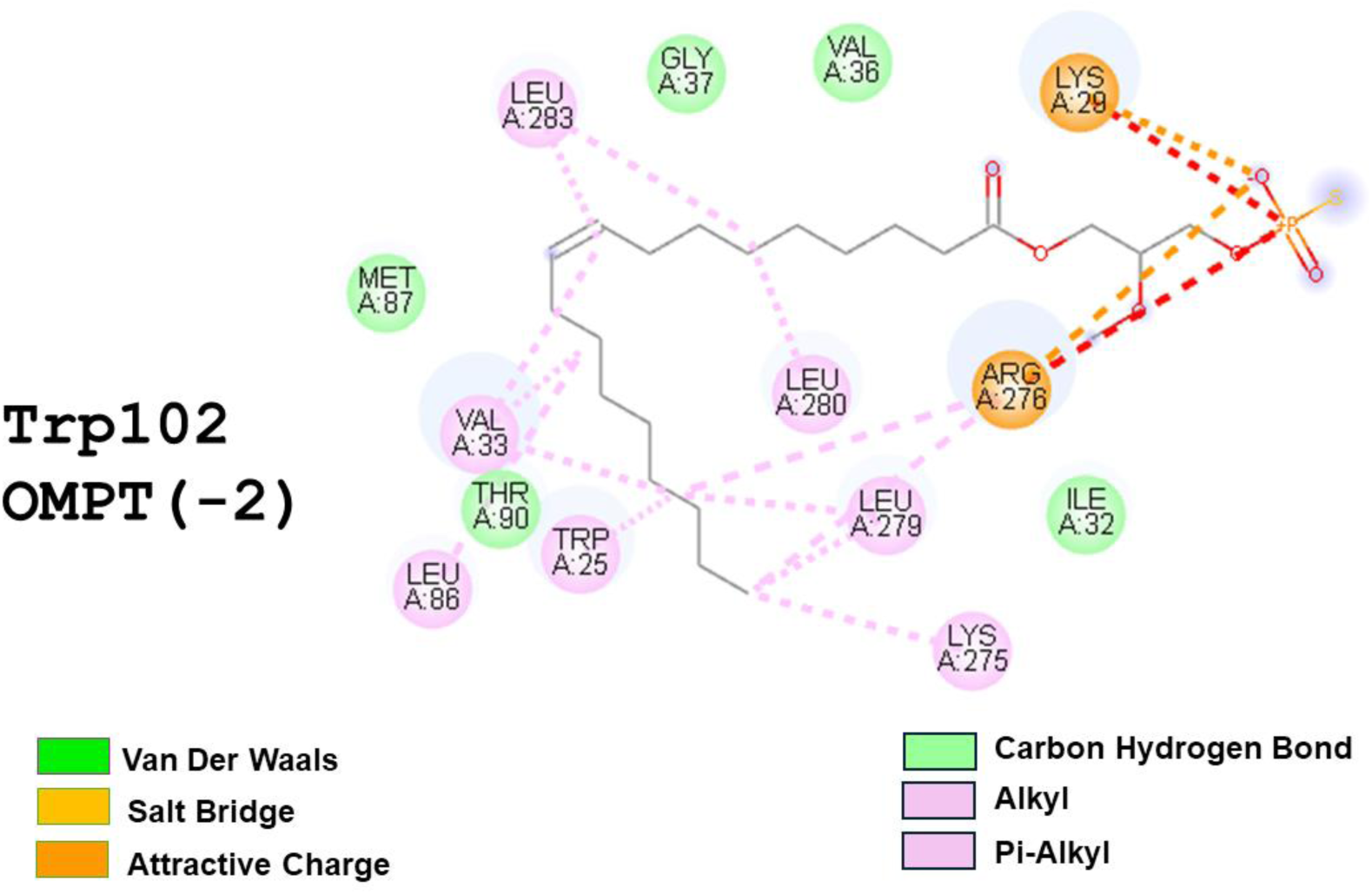
The 2D diagram shows interactions of the LPA_3_ (Trp102) structure with LPA or OMPT. In the individual images Ligand and Charge are indicated (Discovery Studio Visualizer). Critical amino acids contributing to interactions are shown in circles.

**Supplementary Figure S11.**
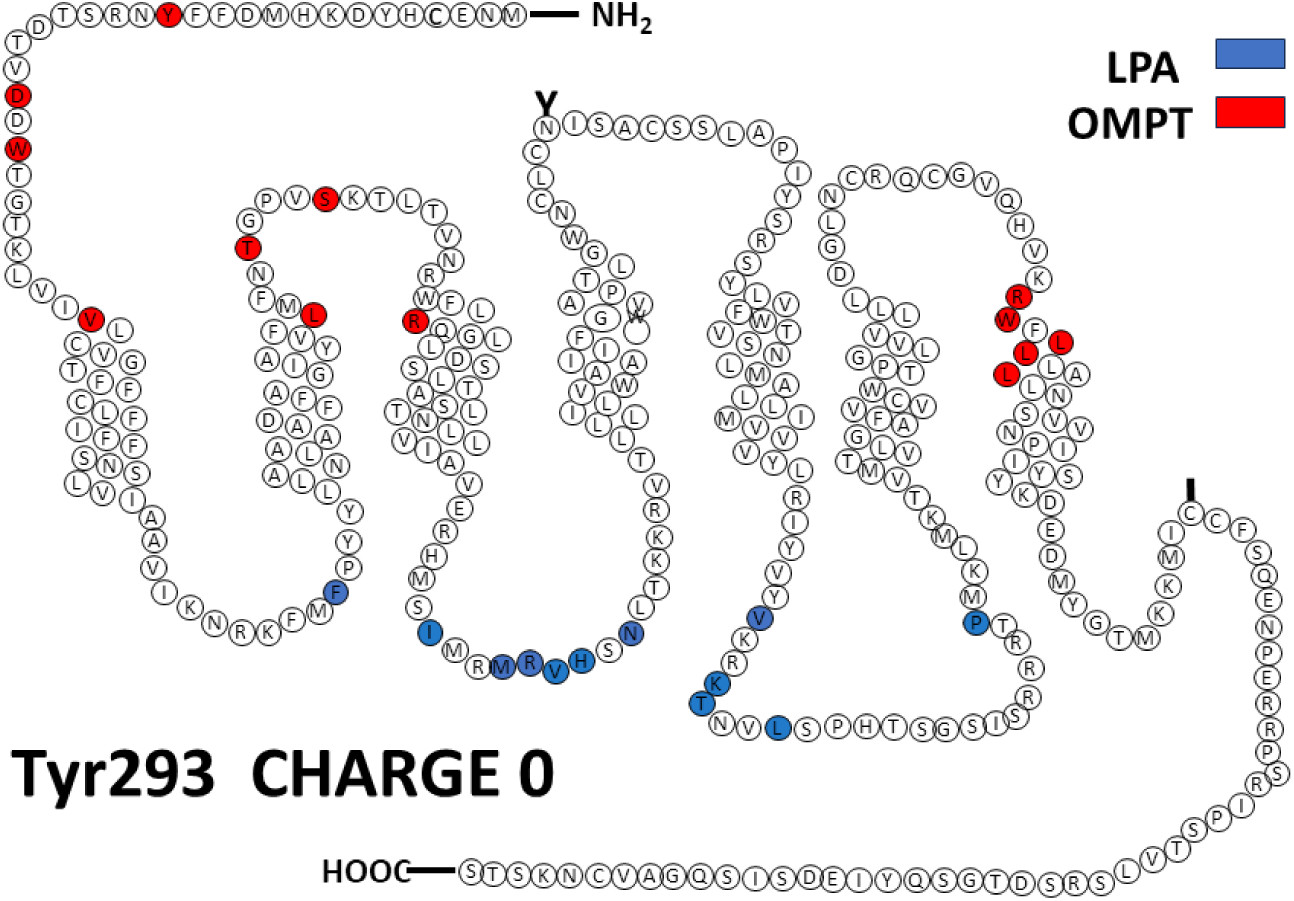
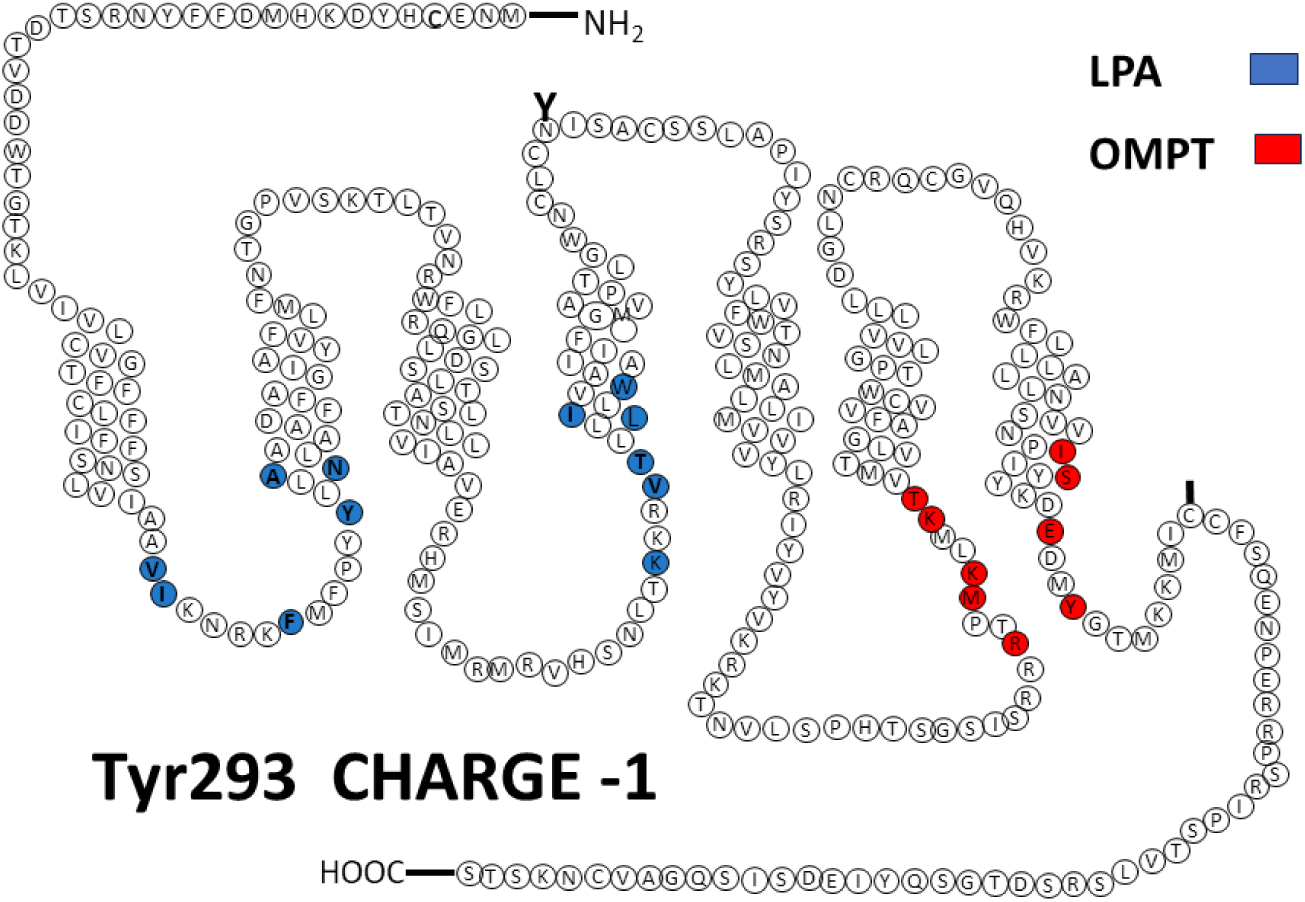
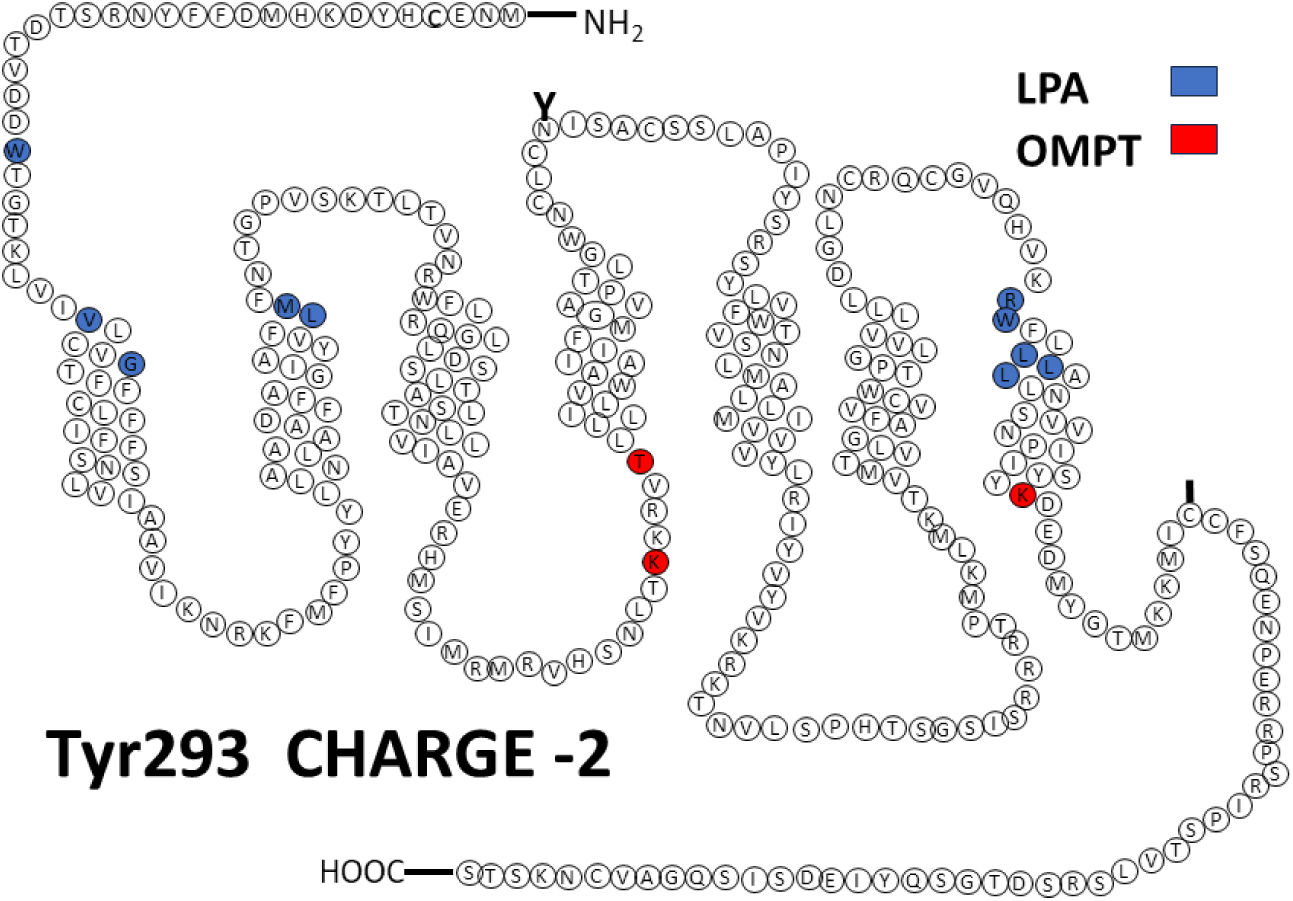
Cartoons showing the LPA_3_ receptor (Tyr293-focused) sites interacting with LPA (blue) and OMPT (red).Charges are indicated in the Figures

**Supplementary Figure S12.**
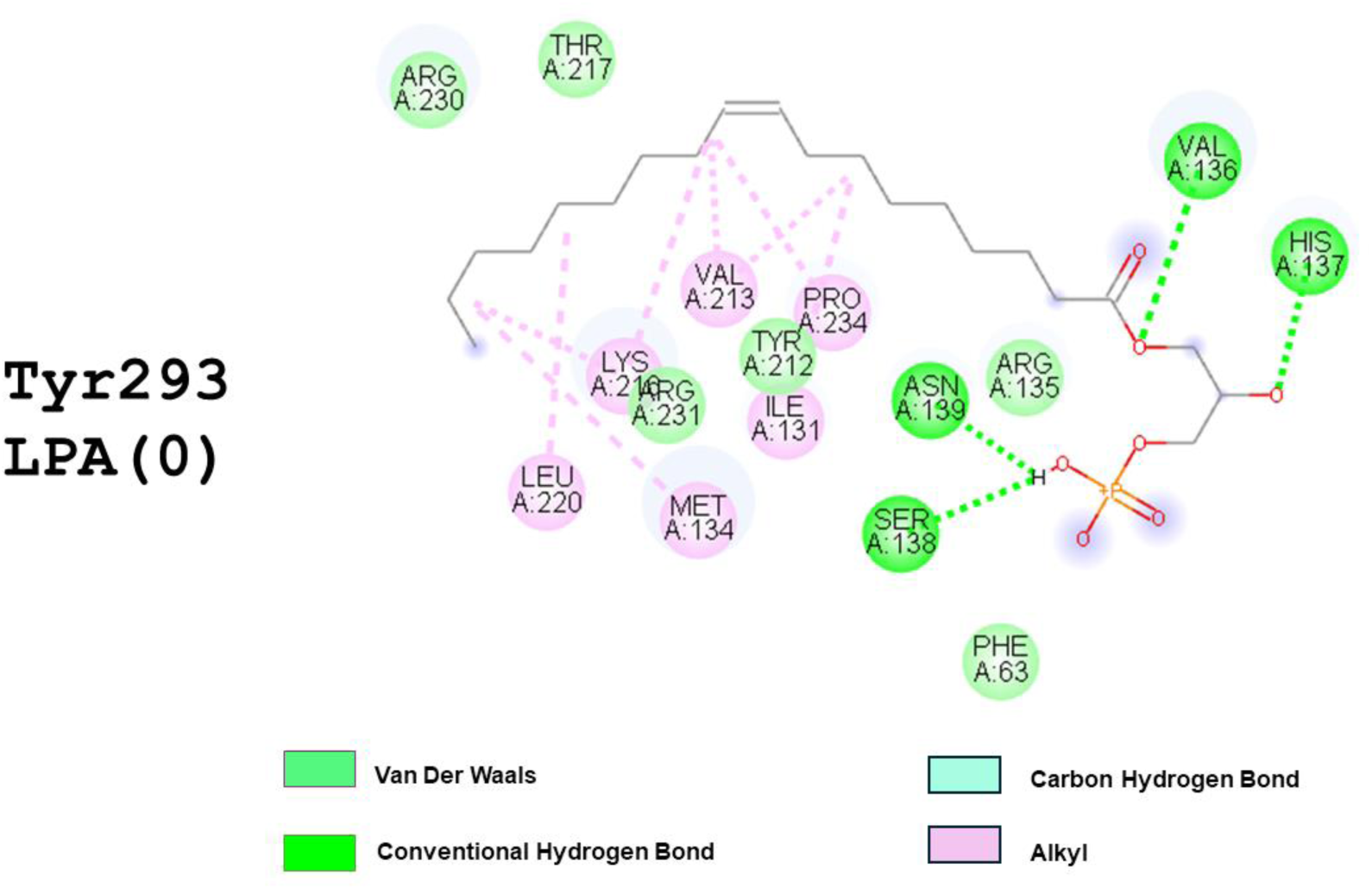
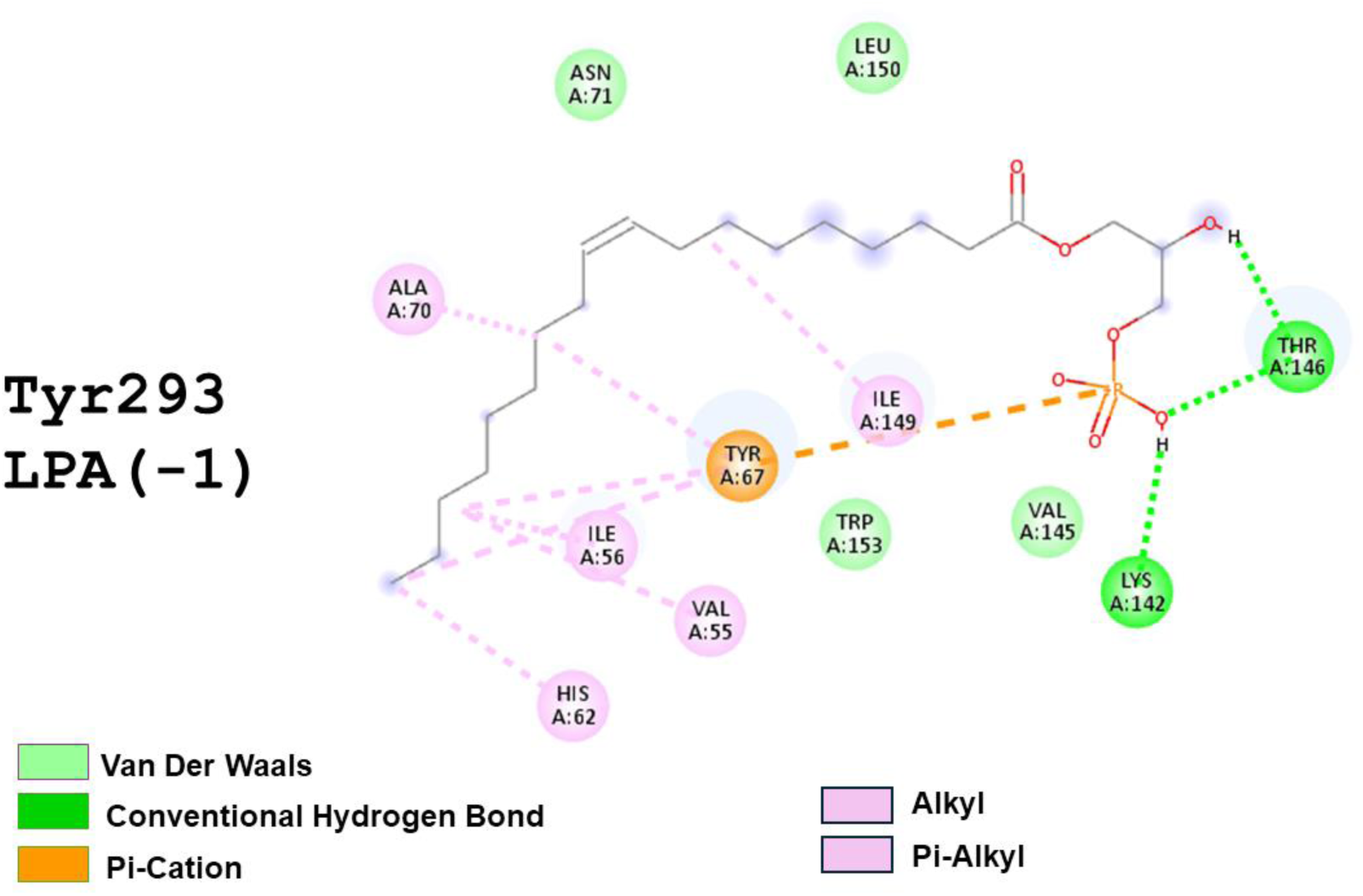
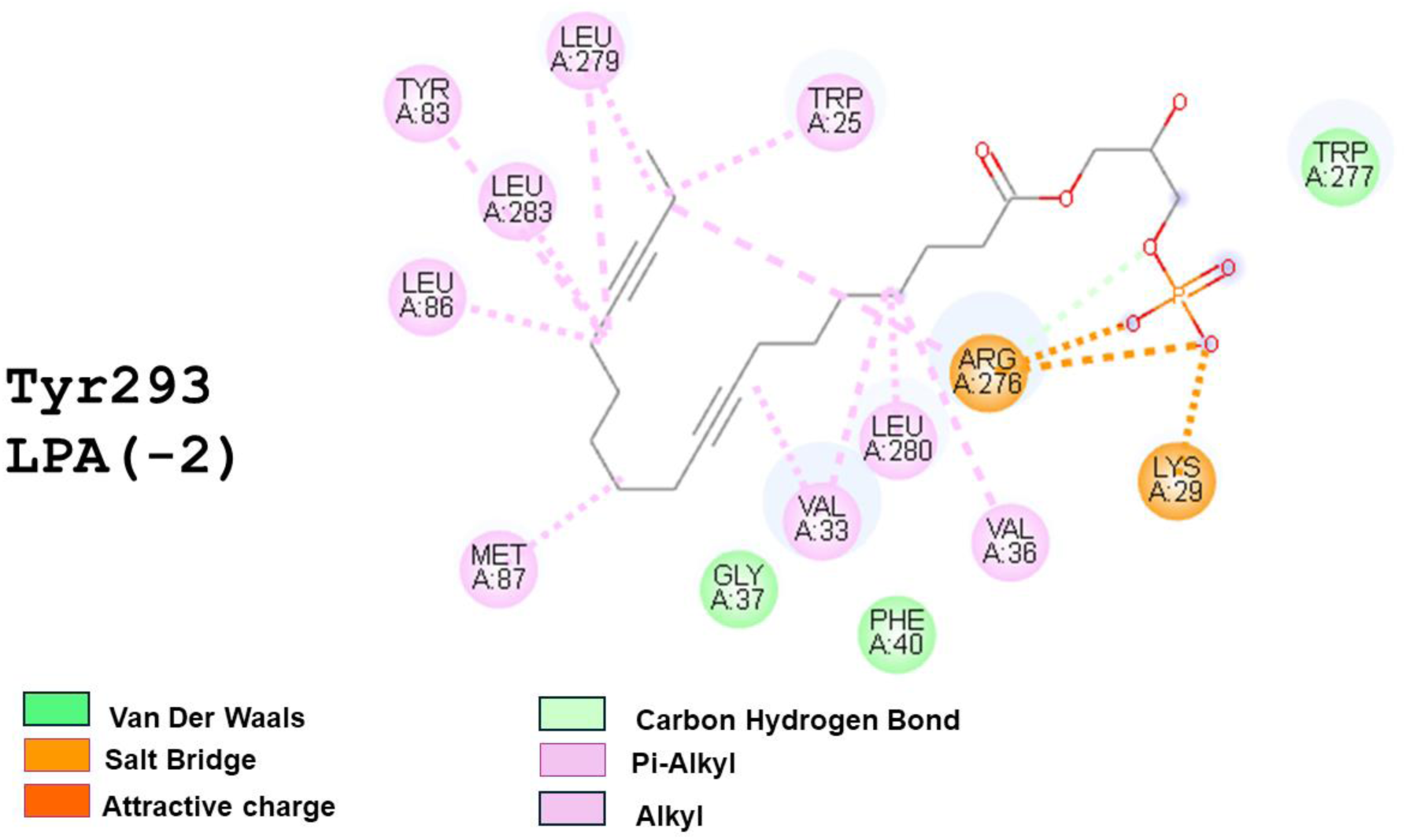
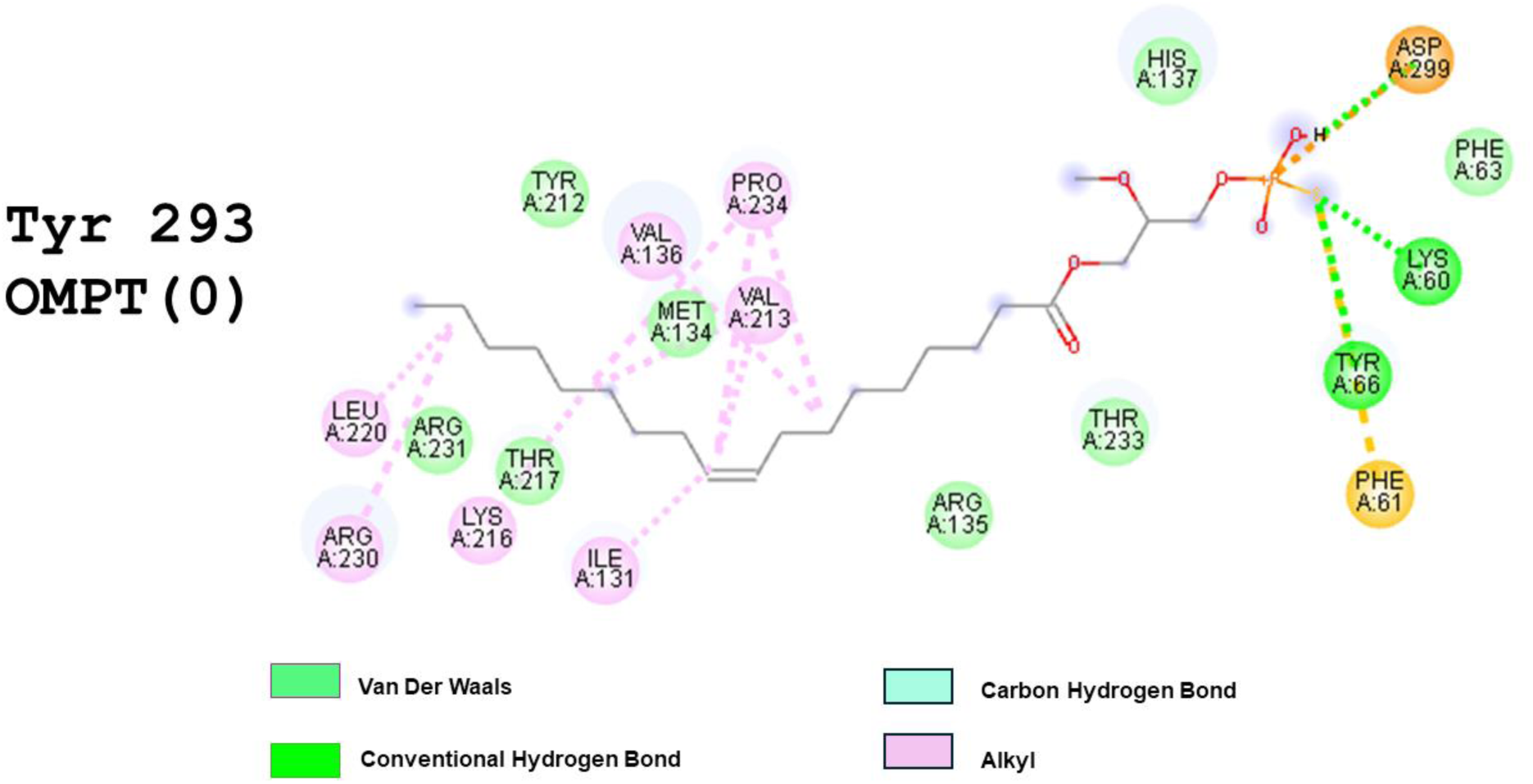
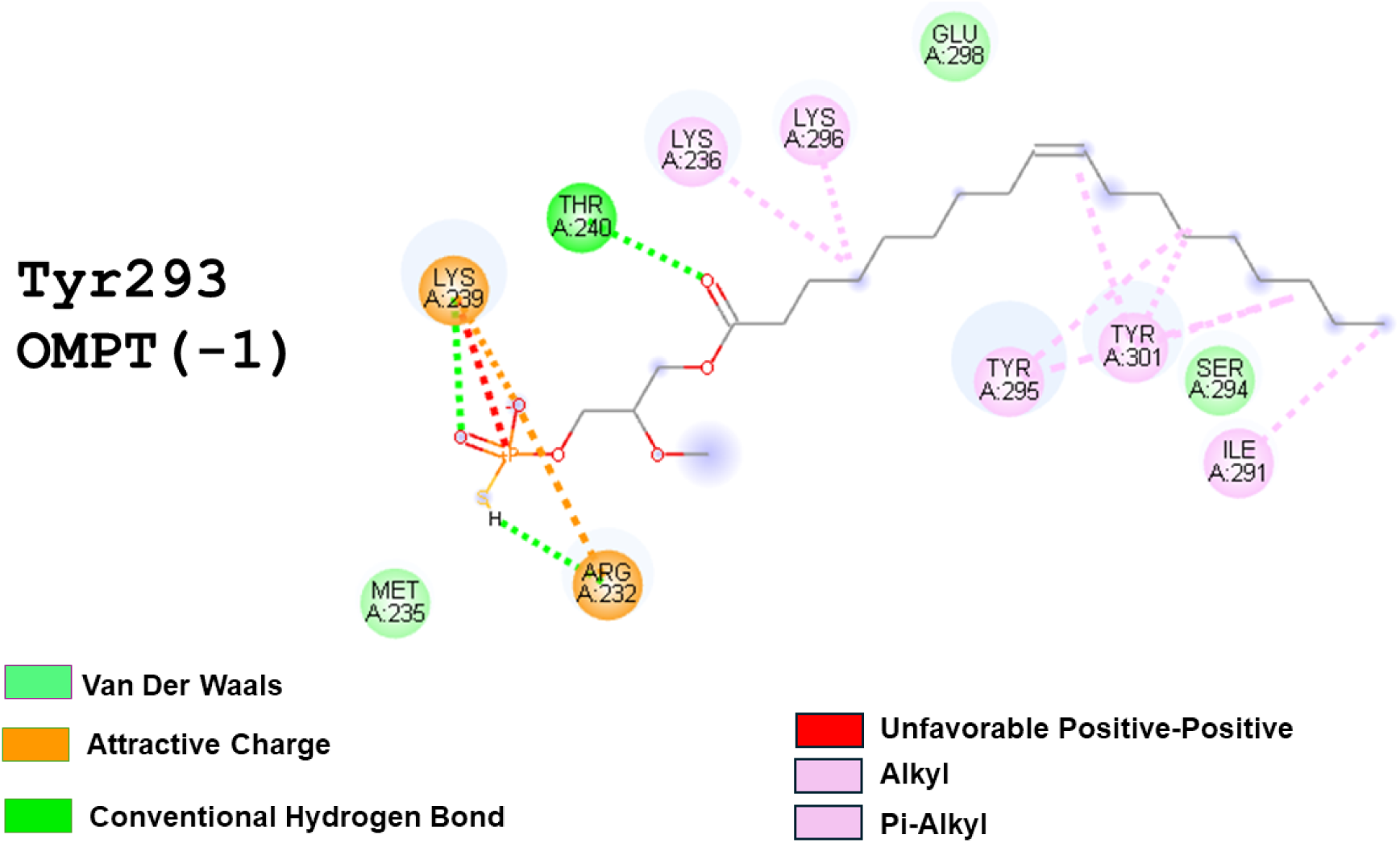
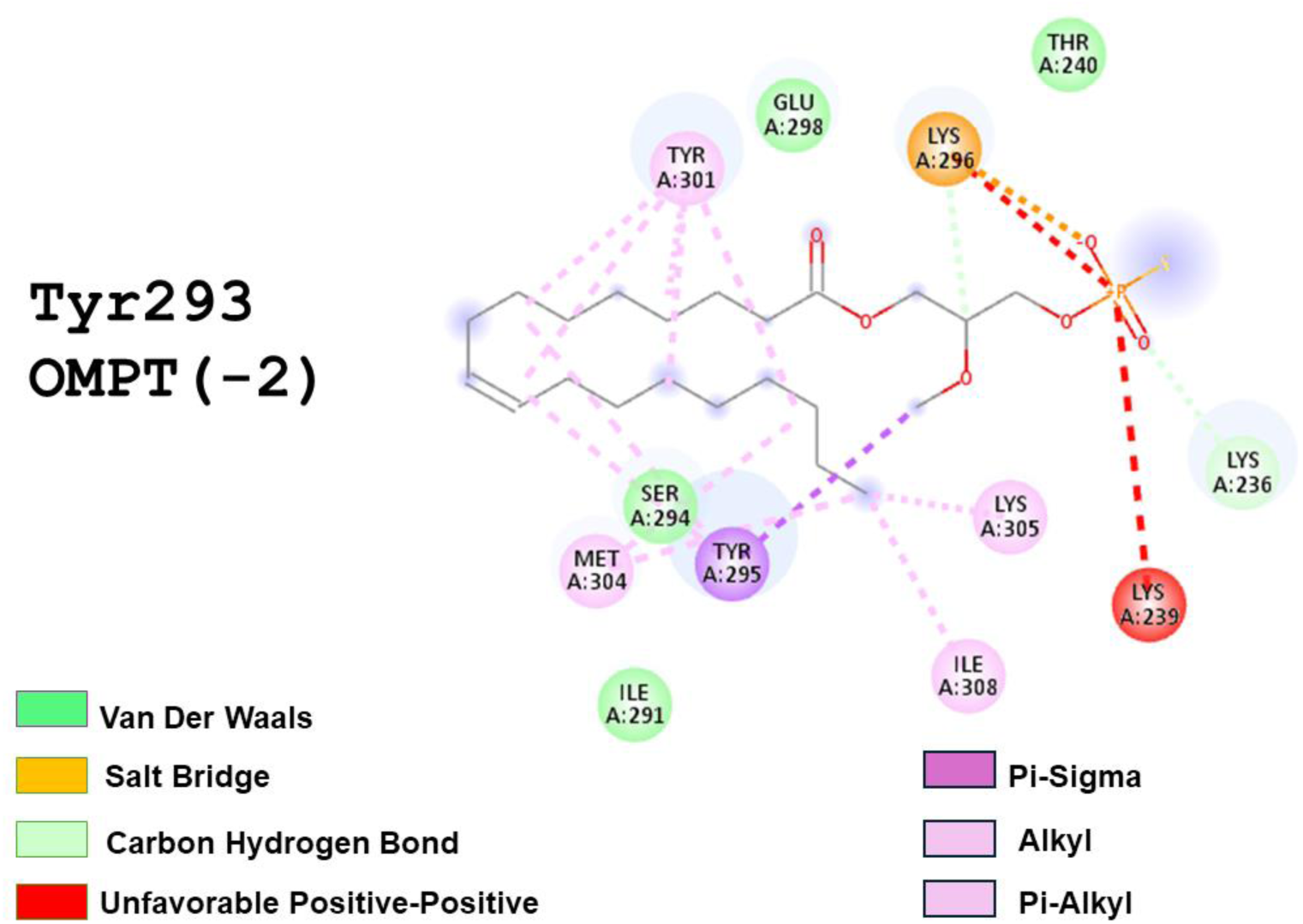
The 2D diagram shows interactions of the LPA_3_ (Tyr293) structure with LPA or OMPT. In the individual images Ligand and Charge are indicated (Discovery Studio Visualizer). Critical amino acids contributing to interactions are shown in circles.

**Supplementary Figure S13.**
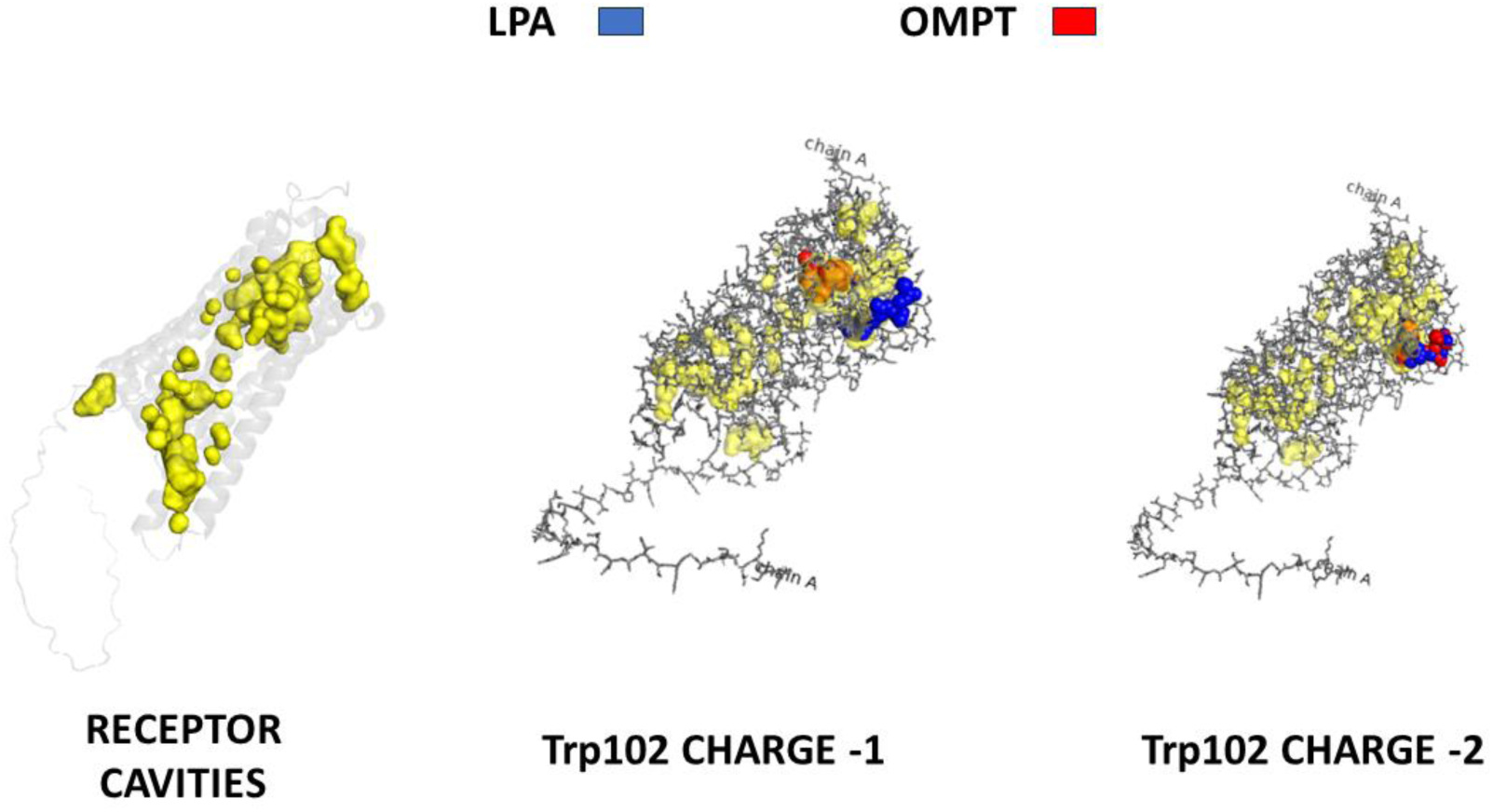
LPA_3_ receptor cavities were evidenced using PyMol. In the left panel the cavities found are indicated in yellow. In the middle (Trp102-focused, Charge-1) and right (Trp102-focused, Charge-1) panel the position of the ligands (LPA, blue, and OMPT, red) are indicated. Change in ligand color is due to merge with the yellow color of the cavity.

---

**Supplementary Table S1.** Structural evaluation of the 3D models generated using the three most frequently used homology modeling servers (in green, the best values).

| Server | MolProbity: Protein Geometry/<br>Ramachandran favored | SAVESv6.0: ERRAT | SAVESv6.0: VERIFY |
| --- | --- | --- | --- |
| SWISS-MODEL | 91.45% | 94.8052 | 33.14% |
| I-TASSER | 78.92% | 83.4302 | 38.81% |
| AlphaFold | 95.18% | 94.3038 | 29.01% |

**Supplementary Table S2.** Blind docking for LPA and OMPT at the unrefined LPA3 receptor structure with a 126 Å3 grid box centered on the protein.

| Ligand | $\Delta G$ (kcal/mol) | Kd ($\mu M$) | Sites of ligand-receptor interaction |
| --- | --- | --- | --- |
| LPA | -3.98 | 1200 | Leu220, Thr217, Arg230, Lys216, Val213, Met134, Ile131, Arg135, Pro234, Asn139, Val136, His137 |
| OMPT | -5.31 | 128 | Leu86, Leu283, Leu279, Leu280, Lys275, Val33, Val36, Thr90, Trp25, Arg276, Trp277 |

